# Mapping Individualized Developmental Imbalance in Youth and Its Association with Psychopathology

**DOI:** 10.64898/2026.05.09.724052

**Authors:** Qingyu Hu, Leo Milecki, Bahram Jafrasteh, Kilian M. Pohl, Amy Kuceyeski, Qingyu Zhao

**Affiliations:** Department of Radiology, Weill Cornell Medicine, New York, NY, USA; School of Electrical and Computer Engineering, Cornell University and Cornell Tech, New York, NY, USA; Department of Psychiatry and Behavioral Sciences, Stanford University, Stanford, CA, USA

## Abstract

The emergence of psychiatric symptoms during adolescence is increasingly hypothesized as arising from maturational imbalance across brain systems. However, this concept largely lacks quantitative grounding, which requires measuring fine-grained imbalance patterns that accurately capture acceleration or delay relative to normative developmental trajectories of brain regions. To address this gap, we leverage predictive normative modeling to learn models that predict chronological age from regional multivariate functional connectivity patterns. We demonstrate that these region-specific models are highly generalizable across independent cohorts and capture greater developmental effects than traditional functional connectivity metrics. From these models, we then derive a region-wise Relative Maturity (RM) index that quantifies individualized, region-specific deviations from normative development. Rigorous cross-cohort and longitudinal evaluations across four datasets show that RM maps are reproducible, subject-specific fingerprints of neurodevelopmental imbalance. These fingerprints are organized along continuous, low-dimensional axes aligned with intrinsic functional gradients and can predict dimensions of psychopathological vulnerability. Together, our findings establish RM as a robust, sensitive, and generalizable framework for quantifying individual vulnerability to psychopathology through system-level patterns of developmental imbalance.

## Introduction

Adolescence represents a critical period of neurodevelopment, characterized by a profound reorganization of the functional connectome underlying the maturation of higher-order cognition, affect regulation, and social behavior [1–4]. This period of heightened plasticity coincides with the peak onset of psychiatric symptoms [5–7], motivating the long-standing hypothesis that psychopathology reflects disruptions in normative brain development (as captured by resting-state fMRI) [8–10]. One influential theoretical perspective is the Dual Systems and Maturational Imbalance models [11–13], which posit that the asynchronous maturation between the faster-developing subcortical socioemotional circuits and the slower, more sustained development of prefrontal cognitive control networks is a risk factor for psychopathology. While conceptually powerful, this framework inherently reduces developmental imbalance to interactions between two specific circuits, whereas cortical maturation unfolds along continuous functional hierarchies spanning the entire cortex [14–17]. Developmental imbalance may therefore be spatially distributed and hierarchically organized, rather than confined to a binary circuit contrast [18].

Empirically capturing such fine-grained imbalance remains challenging. Prevailing approaches to studying functional brain development with resting-state fMRI typically average high-dimensional functional connectivity (FC) into region-level summary metrics [15, 16, 19] and contrast their age-related trajectories across regions. However, such averaging often lacks sensitivity to subtle changes in distributed connectivity patterns during adolescence [20, 21]. A potentially more sensitive approach is prediction-based normative modeling, which uses a machine learning model to predict chronological age based on the whole-brain FC [24]. The predicted age then quantifies an individual’s overall position along a normative developmental trajectory. Recent studies [22, 23] extend this framework to regional analysis by predicting age from the FC of each region to the rest of the brain. Thus, comparing model predictions across regions provides a principled foundation for quantifying region-specific developmental imbalance [25–27].

However, it is unknown whether developmental imbalance quantified by the regional age-prediction models reflects reproducible neurobiological signatures of individuals or is contingent on the cohort used to train the models [28]. Furthermore, it is also unclear whether patterns of developmental imbalance are randomly distributed across the cortex or constrained by intrinsic functional hierarchies. Finally, it remains to be determined whether patterns of developmental imbalance can serve as neurobiological markers that predict psychopathological dimensions at the individual level [29, 30].

To address these three questions, we first establish that regional age-prediction models provide a sensitive and generalizable basis for characterizing regional FC development. Based on this observation, we then derive region-specific relative maturity (RM) as the deviation of predicted development relative to age-matched peers. Finally, we rigorously evaluate the reproducibility, longitudinal stability, and biological organization of RM across three independent cross-sectional cohorts and a large external longitudinal sample [31–35]. We demonstrate that RM exhibits strong cross-cohort invariance and robustness to training data, while preserving a stable, individual-specific neurodevelopmental “fingerprint.” Importantly, this fingerprint is not randomly distributed across the cortex but systematically organized along the intrinsic functional hierarchy of the brain [17]. Using canonical correlation analysis, we demonstrate that these hierarchy-aligned developmental imbalance patterns predict dissociable dimensions of psychopathological vulnerability at the individual level. Together, these findings establish RM as a reproducible, hierarchically organized, and clinically relevant neurodevelopmental marker that refines developmental imbalance theory from coarse circuit contrasts to region-specific metrics.

## Results

### Generalization of Regional Age Prediction Accuracy Across Cohorts

We first evaluated to what extent region-specific FC profiles could serve as reliable markers of local developmental maturity by analyzing resting-state functional MRI from three independent, large-scale adolescent datasets spanning childhood through young adulthood (see Supplementary Fig. 1): HCP-D [31] (*n* = 509; 271 females; age 8–21 years), PNC [33, 36] (*n* = 1,159; 623 females; age 8–23 years), and HBN [32] (*n* = 1,159; 472 females; age 5–21 years) [31–34, 36]. For each scan, whole-brain FC was computed among 400 cortical regions based on the Schaefer 400 atlas (Fig. 1, see Methods) [37]. Next, for each region, we trained a kernel ridge regression model [38, 39] to predict chronological age from the region’s FC to all other regions (Fig. 1, see Methods for confound control). Model accuracy was evaluated by computing Pearson’s *r* between predicted and true chronological age based on within-dataset 10-fold cross-validation and cross-dataset generalization, in which models trained on one dataset were tested on another.

**Fig. 1.**
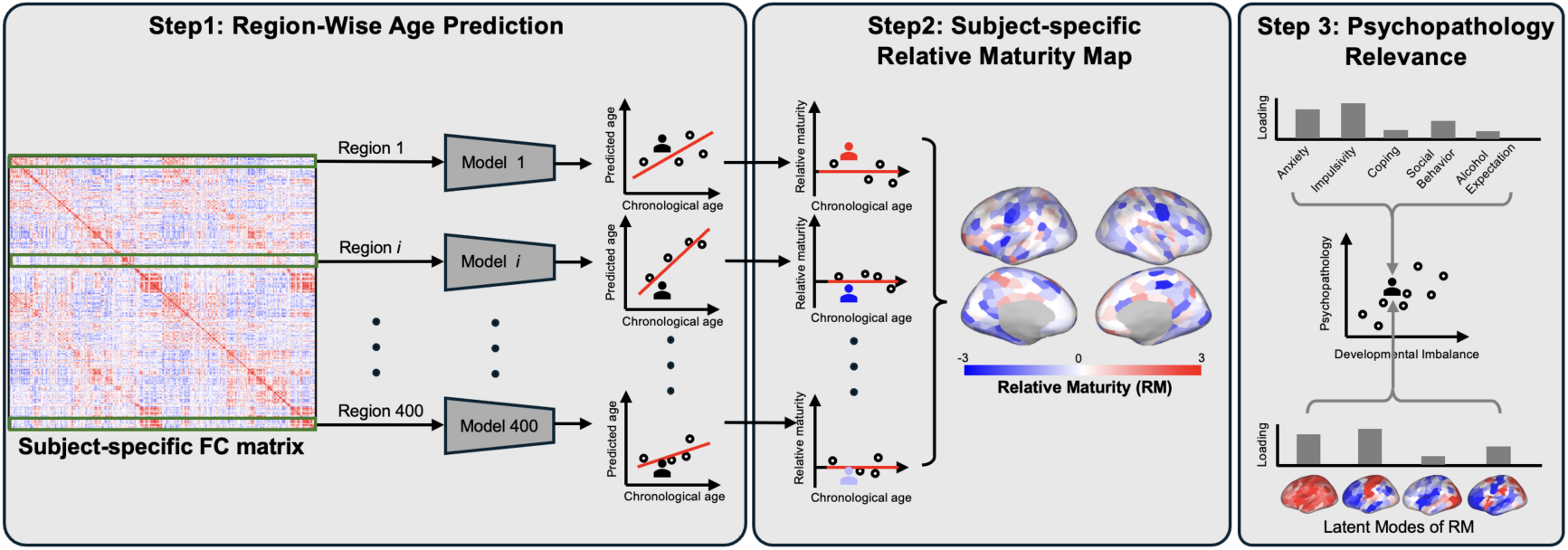
Computational framework for mapping regional relative maturity. The workflow comprises three steps: (1) Region-wise age prediction: Functional connectivity (FC) profiles for each of the 400 cortical regions are extracted and used to train independent, region-specific machine learning models to predict chronological age. (2) Subject-specific Relative Maturity (RM) mapping: The RM index is computed for each region based on the standardized deviation between the predicted and actual age. These regional indices are then combined into an individualized whole-brain map, where red indicates accelerated maturation (positive RM) and blue indicates delayed maturation (negative RM). (3) Psychopathology relevance: Latent spatial variation modes are derived from the RM maps and linked to multidimensional psychopathological scales in individuals.

Regional prediction accuracy based on within-dataset cross-validation ranged from modest to strong correlation between predicted and chronological age (*r* = 0.15–0.70, Fig. 2a). All regions showed statistically significant age prediction after correction for multiple comparisons (Bonferroni-corrected *p* < 0.05). Among the three cohorts, overall prediction accuracy was highest in the HCP-D cohort, with a median regional correlation of *r* = 0.54. When evaluating models trained on sex-balanced cohorts separately in males and females, prediction accuracy remained similar (Supplementary Fig. 2).

**Fig. 2.**
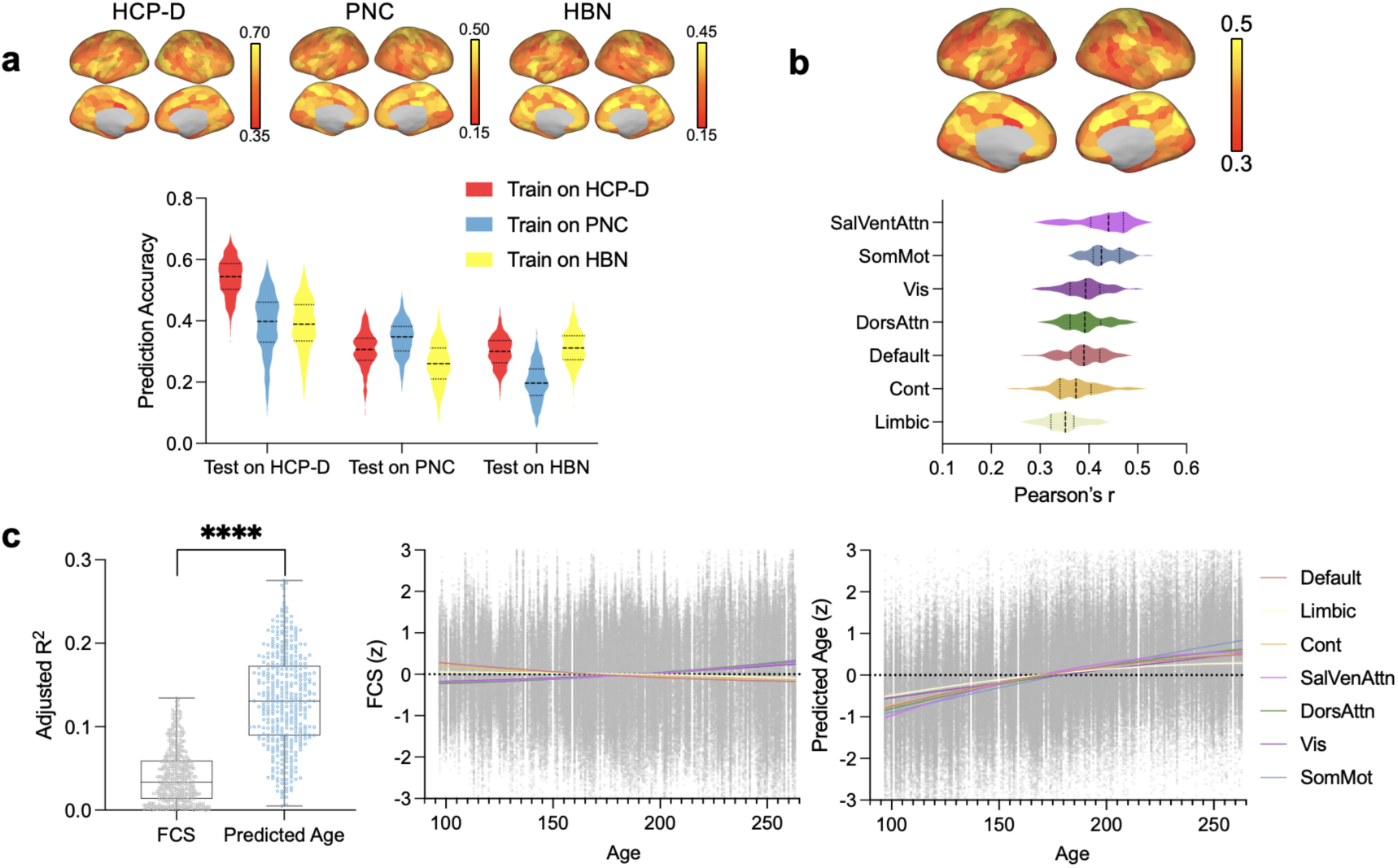
Regional age prediction establishes a robust and sensitive index of local developmental maturity. **a,** Regional age prediction accuracy is highly reproducible across independent datasets. Brain maps (top) display the spatial distribution of within-dataset cross-validated prediction accuracy (Pearson’s *r*) across the 400 cortical regions for the HCP-D, PNC, and HBN cohorts. Violin plots (bottom) illustrate the distribution of regional prediction accuracies for both within- and cross-dataset prediction tasks, indicating robust generalizability. **b,** Predictive performance is spatially heterogeneous and varies by functional topology. The average accuracy map over the three cohorts (top) reveals that prediction accuracy differs by networks (bottom), with highest accuracy in the Salience/Ventral Attention network (median *r* = 0.44) and the Somatomotor network (median *r* = 0.43). **c,** When conceptualized as a **“**maturity index”, the predicted age exhibits superior sensitivity to development compared to mean functional connectivity strength (FCS). Out-of-sample evaluation results on the HCP-D cohort using prediction models trained on the PNC cohort is used to evaluate the sensitivity to development. Left: On the HCP-D dataset, the regional predicted age captures significantly greater age-related variance (adjusted *R*²) than FCS across all 400 regions (paired t-test, *p* < 0.0001). Middle: FCS shows weak or near-flat age trajectories in 7 functional networks. Right: the predicted age exhibits clear increases across all networks, reflecting robust developmental sensitivity.

Visual inspection confirmed that the spatial distribution of regional accuracy was reproducible across all three datasets (Fig. 2a, Supplementary Fig. 3), which was further supported by moderate, significant spatial correlations between regional accuracy scores (*r* = 0.45–0.61, Supplementary Fig. 4). Notably, prediction accuracy remained robust not only based on within-dataset cross-validation but also in stringent cross-dataset generalization settings: when models were trained on a completely independent cohort, median regional accuracy remained stable, ranging from *r* = 0.19 to 0.39 across datasets (Fig. 2b), and the spatial correlation between regional accuracy scores ranged from *r* = 0.33 to 0.73 (Supplementary Fig. 4). Given the high consistency of regional prediction accuracy across the three datasets, we averaged the Pearson’s *r* over the three datasets (Fig. 2b), which revealed that the predictive accuracy was spatially heterogeneous across the 400 regions based on Yeo’s 7–network partition [40]: regions in the Salience/Ventral Attention network exhibited the highest accuracy (median *r* = 0.44), followed closely by the Somatomotor network (median *r* = 0.43). In contrast, the Limbic and Control networks showed weaker neurodevelopment in their FC profiles (median *r* = 0.35 and 0.37, respectively). The Visual, Dorsal Attention, and Default Mode networks displayed moderate predictive performance (median *r* = 0.39).

### Predicted Age as a Sensitive Index of Regional Development

Similar to the brain age prediction framework [24], when a trained model is applied to an unseen individual, the predicted age can be conceptualized as a “maturity index” that quantifies the position of that individual along a normative developmental trajectory. Next, we demonstrate that these region-specific age predictions reveal more pronounced developmental effects than Functional Connectivity Strength (FCS), a commonly used metric defined as the average of a region’s FC with other regions [15, 16]. We first trained the prediction models on the PNC cohort and then applied them to the HCP-D cohort to derive predicted age for each region, ensuring that the models had no access to age information from the HCP-D cohort (see Supplementary Fig. 5 for other training and testing settings). Because the predicted age and FCS differed in scale and distribution, we standardized each metric into a ***z*-score** (using values across all subjects and all regions). Then, for each region, we modeled the relationship between each metric and chronological age using ordinary least squares (OLS) regression (see Methods). To capture potential nonlinear developmental trajectories, age was modeled using both linear and quadratic terms, with sex and head motion included as covariates to control for confounding effects. The extent to which each metric captured developmental effects was quantified using the adjusted *R*^2^ of the OLS model (see Methods).

Based on the OLS regression of FCS, 200 out of the 400 regions showed little to no developmental trend (Bonferroni-corrected *p* > 0.05). In contrast, the predicted age had significantly higher adjusted *R*^2^ across all 400 regions (paired t-test *p* < 0.001, Fig. 2c, left), and 384 out of 400 regions showed significant developmental trends (Bonferroni-corrected *p* < 0.05). The OLS regression curve fitted on regions within each network (Fig. 2c) revealed that model predictions captured pronounced developmental effects across all seven functional networks (adjusted *R*^2^ = 0.05–0.21, Fig. 2c, right), while FCS showed weak or flat trajectories (adjusted *R*^2^ = 0.002–0.05, middle). Particularly, the FCS of the Limbic network showed no significant age-related changes (adjusted *R*^2^ = 0.002), while the age prediction framework detected modest but significant developmental effects in this network. These results validate that region-specific predictive modeling was capable of capturing more comprehensive maturational patterns that are obscured by traditional metrics like FCS.

### Relative Maturity as a Robust Biological Fingerprint

For each region, we fit a linear regression between predicted age and chronological age across the population. The residual captures the extent to which a region’s FC appears developmentally advanced or delayed relative to peers of the same chronological age [22, 41] (Fig. 1, Step 2). We standardized these residuals for each region by *z*-scoring across individuals within the cohort [25] to remove heteroscedasticity in region-specific predictions. We refer to this normalized residual as the Relative Maturity (RM) index. Note, RM is different from prediction error (difference between predicted and chronological age), which often reflects an age-dependent bias (i.e., regression to the mean).

A critical prerequisite for establishing RM as a clinical correlate is its stability: a robust biological fingerprint should reflect the individual’s intrinsic physiology rather than the specific characteristics of the reference cohort used to train the model [28]. To test this, we examined whether the spatial pattern of RM for a given individual remained consistent when derived from models trained on different cohorts. For each individual in the HCP-D cohort, we generated three RM maps: one derived from a model trained within HCP-D using cross-validation, and two derived from models trained on external datasets (PNC and HBN). In all cases, RM values of each region were normalized based solely on the distribution of all HCP-D subjects (see Methods). Similarly, within-dataset and cross-dataset RM maps were derived for each PNC and HBN subject.

Qualitative inspections of individual RM maps confirmed that RM maps were reproducible across training datasets (Fig. 3). As illustrated by three representative subjects, spatial patterns of accelerated (red) or delayed (blue) development were largely preserved, regardless of whether the underlying model was trained on HCP-D, PNC, or HBN data. Quantitative analysis further confirmed this stability. For each subject in HCP-D, PNC, and HBN, we computed pairwise spatial correlations between RM maps derived from models trained on different cohorts. This analysis revealed consistent, robust cross-model agreement across all training and testing combinations, with median spatial correlations consistently approximating Pearson’s *r* ≈ 0.40 (Fig. 4a). This suggests the estimated RM topography captures stable, individual-specific neurodevelopmental fingerprints.

**Fig. 3.**
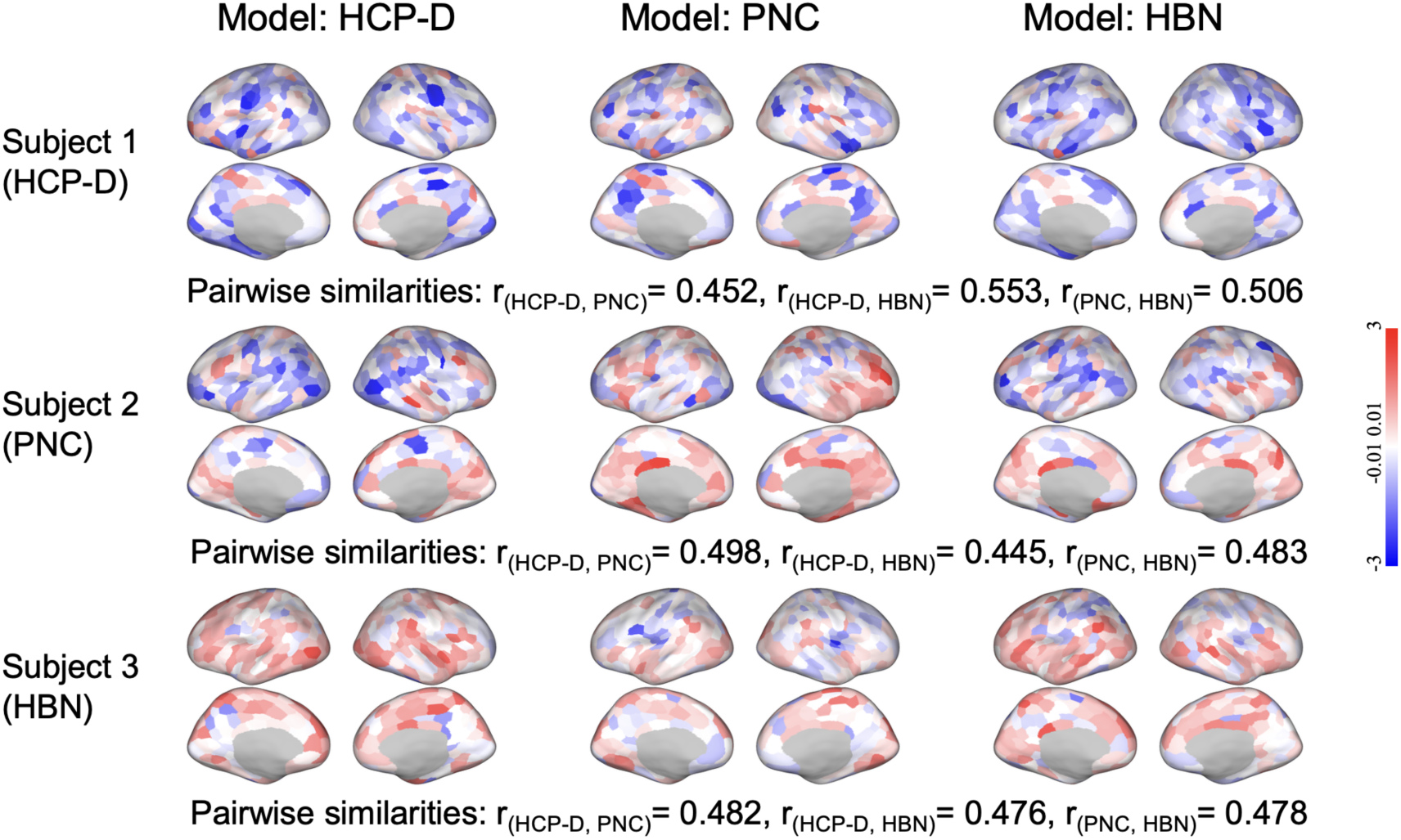
Samples of individualized Relative Maturity (RM) maps derived from models trained on different cohorts. We derived RM maps for three representative subjects (rows) from the HCP-D, PNC, and HBN cohorts using models trained on three different datasets (columns). Despite differences in the underlying training data, the spatial distribution of accelerated (red) and delayed (blue) cortical maturation is highly conserved within each individual. Pairwise spatial similarities (*r*) between the model outputs confirm this stability, demonstrating that the derived RM maps robustly capture reliable, subject-specific patterns of developmental asynchrony.

**Fig. 4.**
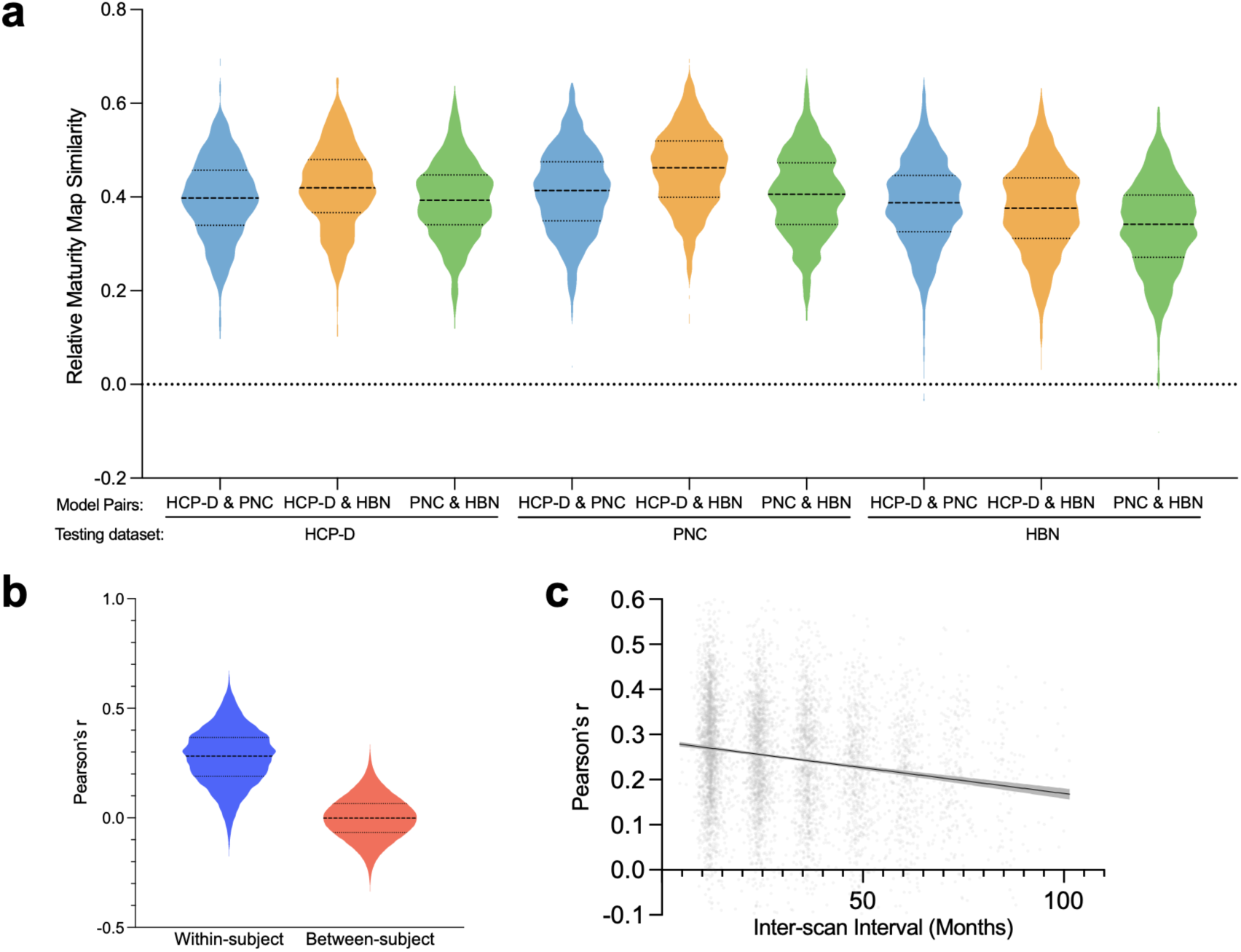
Robustness, individual specificity, and longitudinal sensitivity of Relative Maturity (RM) maps. **a,** Individualized Relative Maturity (RM) maps demonstrate high cross-model robustness. Violin plots show spatial correlations (Pearson’s r) between RM maps generated using models trained on different cohorts (HCP-D, PNC, and HBN) when applied to the same testing individuals. Median spatial correlations consistently approximate *r* ≈ 0.40, indicating that RM topography is robust to the choice of training data. **b,** The RM index exhibits strong individual specificity. Violin plots compare the spatial correlation of RM maps between two longitudinal scans of the same individual (blue) versus the spatial correlation between scans of two different, randomly selected individuals (red). The within-subject similarity is significantly higher (unpaired t-test, *p* < 0.0001), establishing a reliable, individual-specific developmental fingerprint. **c,** Within-subject RM topography is sensitive to longitudinal developmental change. The scatter plot illustrates the relationship between the time interval (age difference) between two longitudinal scans of the same individual and the spatial correlation of the two corresponding RM maps. The linear regression fit (solid line) and its 95% confidence interval (shaded envelope) reveals a significant time-dependent decay in spatial similarity (*r* = -0.17, *p* < 0.0001), demonstrating that the RM index captures dynamic developmental trajectories.

Having demonstrated that RM estimates were largely invariant to the training dataset, we next tested whether RM was stable over time within individuals and whether it captured longitudinal developmental change. We tested these properties on a large-scale external longitudinal dataset from the NCANDA study [35], comprising 798 participants aged 12 to 21 years who had no structural anomaly at baseline and were scanned annually. We trained the age prediction models on the combined cohort of HCP-D, PNC, and HBN (age range 5–22 years) and derived the RM maps for all 3,199 rs-fMRI scans of NCANDA collected across the 12–22 years age range (4.01±1.98 annual scans per participant).

First, we assessed the individual specificity of RM by quantifying the spatial correlation between RM maps from two randomly selected longitudinal time points from the same individual. As shown in the violin plots (Fig. 4b), the distribution of within-subject correlations significantly exceeded correlations between RM maps from two different randomly selected individuals (*p* < 0.0001, unpaired t-test). This validates that RM preserves a stable, individual-specific “identity” of neurodevelopment distinct from group-level variance.

Second, we tested whether RM demonstrated sensitivity to longitudinal change. If RM were merely reflecting static individualized functional topological features, the within-subject similarity would remain constant regardless of the time elapsed between scans. Instead, we observed a significant, time-dependent decay in spatial correlation: as the interval between two longitudinal scans of the same individual increased, the topological similarity of the RM maps significantly decreased (*r* = -0.17, *p* < 0.0001, Fig. 4c).

### Latent Spatial Structure of Relative Maturity Variation

Having established that the RM index is both individually specific and dynamically sensitive to longitudinal change, we next asked whether the spatial pattern of developmental imbalance itself follows an organized principle across the brain. While Fig. 3 reveals substantial heterogeneity in developmental imbalance across regions within each individual and across individuals, this variation exhibits clear spatial autocorrelation, with functionally similar regions tending to mature in synchrony. This distinct spatial coherence suggests that individual developmental deviations are not random, but instead follow a latent spatial principle. To characterize this latent structure at the population level, we aggregated the RM maps of all subjects across the three datasets (HCP-D, PNC, and HBN; total *n* = 2827), each derived using within-dataset model training. We then performed principal component analysis (PCA) on this combined data to extract low-dimensional modes of developmental imbalance.

This analysis revealed a hierarchical organization of RM maps, composed of modes of developmental imbalance patterns. The first principal component (PC1) exhibited uniform loading signs across virtually all cortical regions (Fig. 5a, left), suggesting a “Global Maturation Factor”, a dimension distinguishing individuals who are globally advanced in their neurodevelopment from those who are globally delayed. To validate this interpretation, we computed a whole-brain relative maturity score for each subject, utilizing a separate model predicting age from whole-brain FC features (see Methods). This global metric showed a strong association with PC1 (*r* = 0.64, *p* < 0.0001; Fig. 5a, right) but showed negligible associations with higher-order components (*r* < 0.06, for PC2–5, Supplementary Fig. 6). This selective correspondence confirms that the dominant source of variance in the population (Supplementary Fig. 7) reflects the overall pace of maturation.

**Fig. 5.**
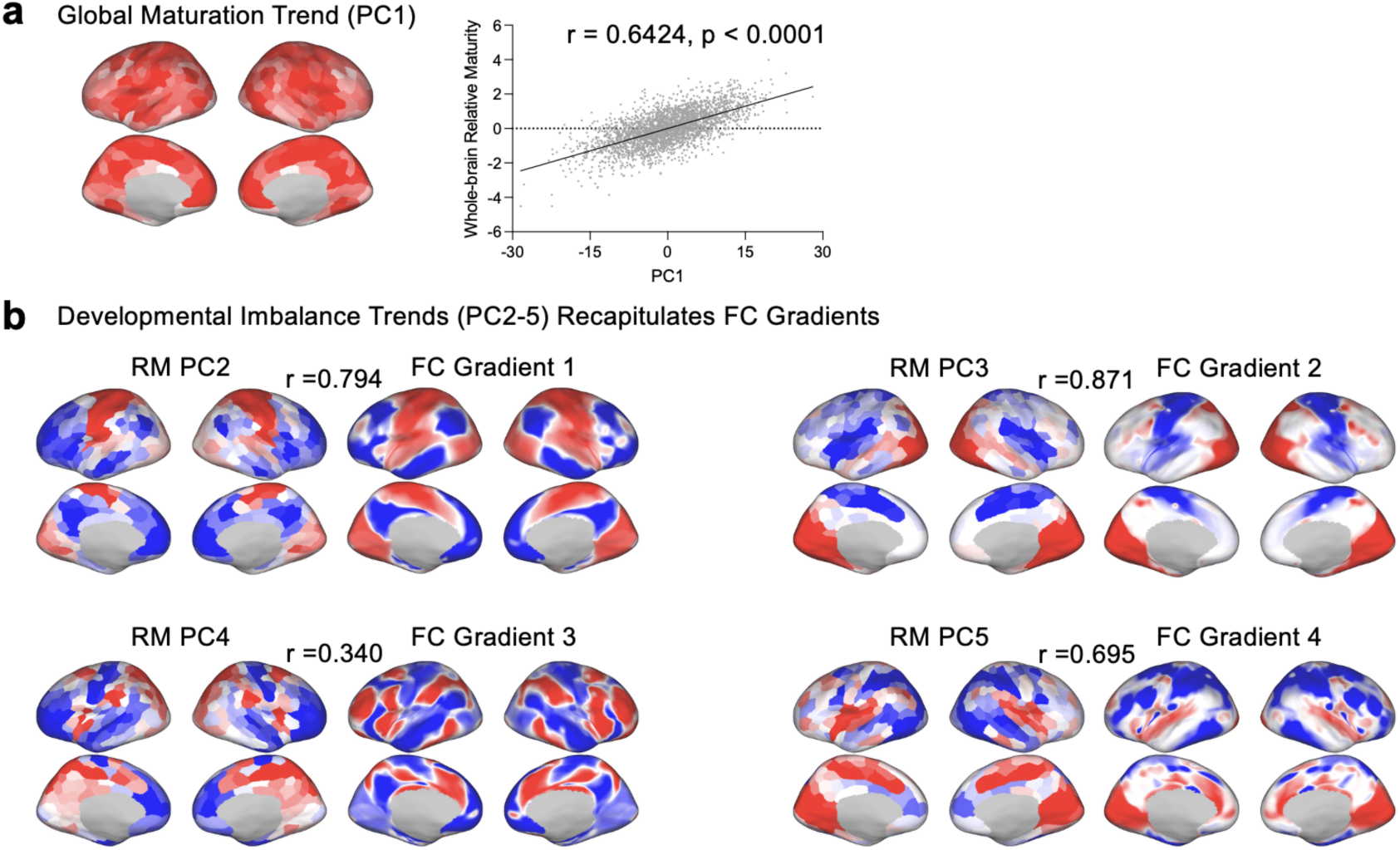
Low-dimensional modes of developmental imbalance recapitulate functional cortical gradients. a, Global maturation trend. Principal Component Analysis (PCA) reveals a dominant first component (PC1) characterized by uniform loading signs across all cortical regions (left). This global factor correlates strongly with subject-specific whole brain relative maturity (*r* = 0.64, *p* < 0.0001), representing a “global pacing” dimension where development is synchronized (globally advanced or delayed) across the cortex. **b, Developmental imbalance trends.** Higher-order principal components (PC2–5) of RM capture specific spatial patterns of developmental imbalance. These imbalance modes (left in each pair) strongly align with the principal gradients of FC (right in each pair). For example, Maturity PC2 tracks the sensorimotor-association axis (FC Gradient 1; *r* = 0.79, *p*_spin_ < 0.0001), while Maturity PC3 recapitulates the visual-somatomotor dissociation (FC Gradient 2; *r* = 0.87, *p*_spin_ < 0.0001). This topographic correspondence indicates that developmental asynchrony is not random but is constrained by the brain’s underlying functional organization.

Beyond the global factor, higher-order components (PC2–5) uncovered common, latent modes of developmental imbalance, where accelerated maturation in specific brain systems is intrinsically coupled with delayed maturation in others. These modes of asynchrony strongly recapitulated the intrinsic functional gradients of the human cortex (Fig. 5b) [17]. Notably, the second principal component (PC2), the dominant mode of developmental imbalance, exhibited a robust spatial alignment with the principal gradient of FC along the Sensorimotor-Association (S-A) axis (*r* = 0.794, *p* < 0.0001, spin test). Similarly, PC3 tracked the second functional gradient along Visual-Somatomotor dissociation (*r* = 0.871, *p* < 0.0001, spin test), and PC4 and PC5 tracked the third and fourth functional gradients (Fig. 5b). These spatial axes of developmental imbalance were successfully replicated in the NCANDA dataset (Supplementary Fig. 8). This tight topographic correspondence indicates that asynchronous individual deviations in neurodevelopment preferentially emerge along the axes that define cortical hierarchy.

### Psychopathological Relevance of Relative Maturity

Lastly, we tested whether RM serves as a biomarker of psychopathology based on a subset of 772 NCANDA participants (*n* = 2,369 scans) who possessed a complete battery of psychopathological and cognitive assessments [42]. Given the strong hierarchical organization of RM revealed previously, we applied canonical correlation analysis (CCA) [43] to relate the PC scores of the RM to 37 psychopathological scales. CCA derived a set of canonical components, each represented by a pair of linear combinations of PC scores and psychopathological scales that were maximally correlated. We systematically varied the number of input PCs (from 1 to 20) to evaluate the out-of-sample correlation in a 10-fold cross-validation. Based on a permutation test, the first canonical component demonstrated significant out-of-sample correlation (Supplementary Fig. 9), which peaked when using 6 PCs (*r* = 0.13, *p* < 0.0001, Fig. 6a). These findings were further replicated in a subset of 645 NCANDA participants with no-to-low alcohol use at baseline (Supplementary Fig. 10a).

**Fig. 6.**
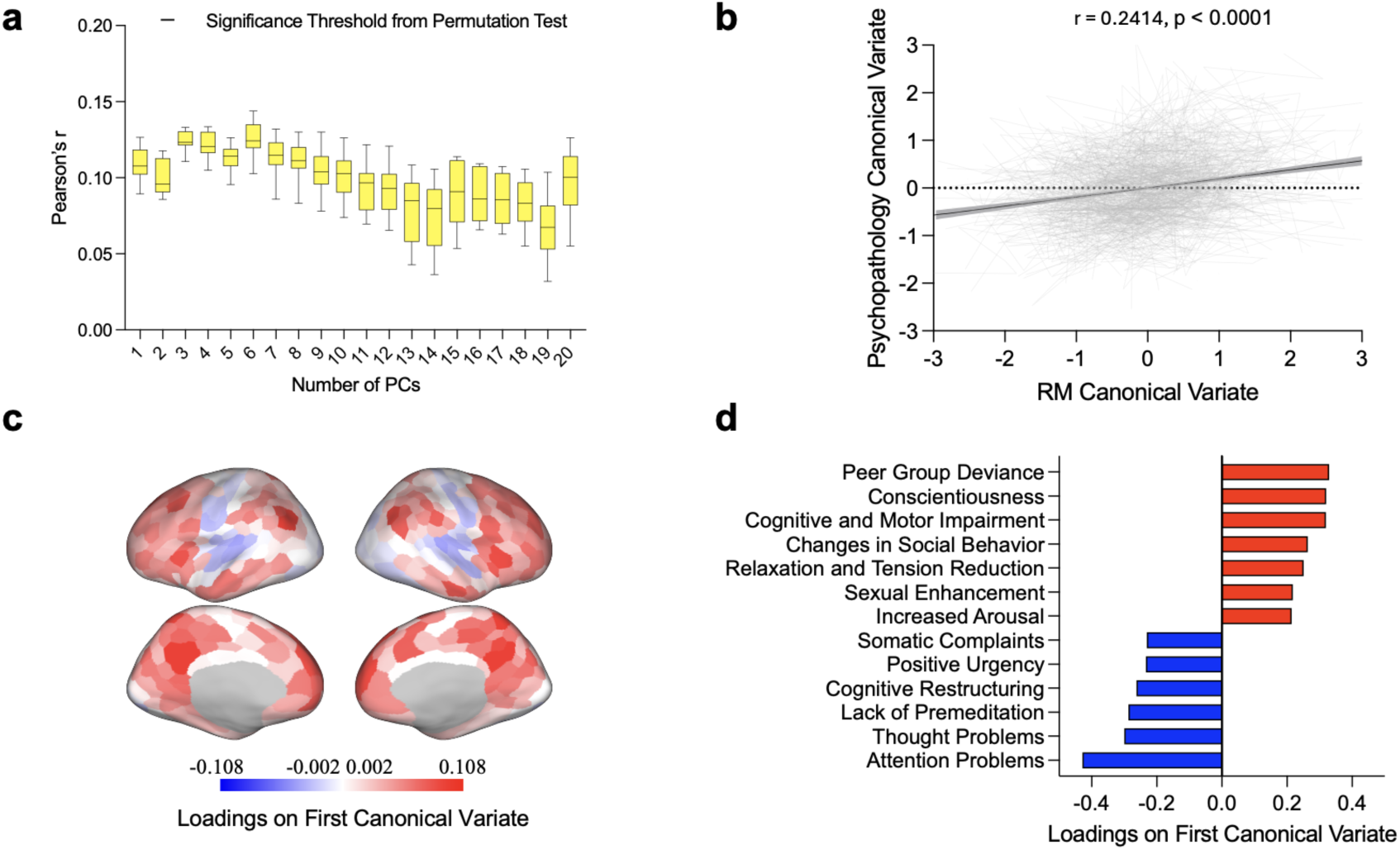
Relative Maturity maps reveal a robust neurodevelopmental axis of psychopathological vulnerability. **a,** Out-of-sample prediction confirms the association between neurodevelopmental imbalance and psychopathological scales. We performed Canonical Correlation Analysis (CCA) to relate RM maps to 37 behavioral measures in 772 NCANDA participants. The model used a varying number of Principal Components (PCs) derived from RM maps (1–20 PCs) as input features. Box plots show the 10-fold cross-validated prediction accuracy (Pearson’s *r*) of the first canonical component. Prediction performance peaks at 6 PCs, indicating stable and robust associations between low-dimensional patterns of developmental imbalance and multidimensional psychopathological profiles. **b,** Canonical variates of RM and psychopathological scales are highly correlated (derived from the CCA model utilizing 6 PCs trained on all NCANDA data). **c,** Based on the CCA model with 6 input PCs trained on all NCANDA data, we visualize canonical loadings of brain regions from the first canonical component to indicate which regions contribute most strongly to the relationship between Relative Maturity (RM) and behavioral measures. The cortical map highlights a specific “imbalance phenotype” that contrasts positive loadings in transmodal association cortices (red) with negative loadings in unimodal sensorimotor and visual regions (blue). **d,** Psychopathology scales (with large canonical loadings > 0.2) contributing most strongly to the RM-psychopathology association.

Next, we examined the canonical loadings of brain regions (see Methods) to interpret which RM patterns contributed most strongly to the significant brain–behavior association. The resulting brain loadings (Fig. 6c) recapitulated the S-A axis (*r* = 0.54, *p* < 0.0001, spin test), the principal gradient of cortical organization [14]. The map was characterized by a divergent pattern: positive loadings in transmodal association cortices (red, such as the Default Mode and Frontoparietal networks) contrasted with negative loadings in unimodal sensorimotor and visual regions (blue). This pattern represents a specific “imbalance phenotype”, a decoupling between the maturation of higher-order cognitive systems and lower-order processing systems. This pattern was not driven by baseline age differences (Supplementary Fig. 11) and persisted when replacing the 6 PCs with 3 to 18 PCs as the input to the CCA model (Supplementary Fig. 12).

To identify which dimensions of psychopathology were associated with this imbalance phenotype, we examined psychopathological scales with large canonical loadings (|*r*| > 0.2). Fig. 6d suggests that faster development in sensorimotor than association cortices was linked to worse dysregulation and internalizing symptoms (Attention Problems, Thought Problems, Somatic Complaints), and higher Impulsivity (Lack of Premeditation and Positive Urgency). Conversely, faster development in association cortices was linked to social and reward-oriented risks, including Peer Group Deviance, Conscientiousness, and a broad spectrum of positive Alcohol Outcome Expectancies, specifically expectations of social facilitation, relaxation, and sexual enhancement.

## Discussion

In this study, we leveraged four large-scale, independent datasets (*n* = 3,625) to establish a computational framework (Fig. 1) for mapping the heterogeneity of adolescent neurodevelopment and to evaluate its biological and psychopathological relevance. Through stringent cross-dataset evaluation and comparison with traditional metrics (Fig. 2), we showed that age prediction based on region-specific machine learning models is a robust, reproducible, and highly sensitive estimator of local functional developmental stages. Based on the prediction, we derived relative maturity maps and showed that they captured stable, subject-specific functional fingerprints of developmental imbalance, while remaining dynamically sensitive to longitudinal change (Fig. 3-4). Notably, the spatial patterns of relative maturity can be decomposed into a global maturation factor and multiple modes of developmental imbalance that are systematically organized along the brain’s principal FC gradients (Fig. 5). Lastly, we revealed that these hierarchy-aligned developmental imbalance patterns were significantly associated with dimensions of psychopathology (Fig. 6). Such a framework provides a biologically grounded basis for understanding heterogeneity in mental health risk and may facilitate more precise identification of individuals at risk based on system-level developmental profiles rather than region-specific abnormalities.

Previous research has demonstrated spatial variability of FC development, with distinct cortical systems exhibiting different developmental trajectories [27, 44, 45]. Inspired by recent advances in region-wise predictive modeling [22, 27], we used age prediction accuracy by machine learning models to quantify the extent to which FC is regionally reorganizing.

Extending the spatial topography of prediction accuracy established in single datasets [27], we demonstrated that this pattern of accuracy is reproducible across three large datasets in both within-dataset and stringent cross-dataset evaluation schemes. Specifically, the Ventral Attention and Somatomotor networks consistently exhibited the highest age prediction accuracy, suggesting that these two networks show the most pronounced functional reorganization during adolescence. This finding aligns with recent evidence identifying the Ventral Attention network as a primary driver of connectome maturation [21], likely reflecting its role as an active transitional hub coordinating large-scale network reconfiguration. Similarly, the Somatomotor network undergoes continued development from childhood to adolescence, marked by significantly increased intrinsic connectivity within the sensorimotor cortex alongside decreased coupling with the Default Mode network [15, 16]. Such concurrent processes of integration and segregation create a regional profile that serves as a potent predictor of biological age.

Indeed, multivariate regional FC profiles provide a more comprehensive characterization of developmental reorganization than traditional univariate metrics. For example, FCS averages connectivity changes across regions [15, 16], such that increases and decreases in connectivity can cancel each other out, potentially obscuring meaningful developmental dynamics. This limitation is particularly evident in the Limbic network, which, during adolescence, shows functional integration with sensorimotor and Frontoparietal networks and functional segregation from the Default Mode network [16]. Such opposing directional shifts result in near-zero change in FCS. In contrast, the multivariate predictive framework captures the distributed patterns of connectivity change, substantially increasing sensitivity to maturation-related effects even in the limbic system.

Given the high sensitivity to age effects, recent studies have used the prediction residuals of machine learning models to quantify RM maps [22, 27], i.e., whether a region showed accelerated or delayed functional development compared to the norm. However, whether the RM map can be stably derived on the regional level with cross-cohort reproducibility has not been studied. Our cross-reference validation revealed that Relative Maturity maps generated from models trained on independent datasets yielded highly concordant individual topographies. This cross-cohort consistency indicates that the specific choice of training data does not substantially alter the estimated RM map, validating the index as a robust biological fingerprint. Furthermore, these individualized patterns were preserved across longitudinal scans while remaining sensitive to time-dependent changes, highlighting the potential of RM maps for tracking longitudinal functional trajectories.

Having established that RM maps capture reliable, individualized biological fingerprints, our study also revealed organizational principles of these unique topographies. Adolescent neurodevelopment is intrinsically heterogeneous [46–50]. Previous research has primarily focused on characterizing developmental trajectories at the population level, showing that different cortical systems mature along distinct timelines [27, 51, 52]. However, it remains unclear whether the resulting patterns of developmental imbalance, captured here by RM maps, also vary systematically across individuals. Recent work using clustering approaches has proposed that individuals’ RM maps might be grouped into categorical subtypes [27]. In contrast, our findings suggest this heterogeneity is better described as continuous axes of variation in a low-dimensional latent space, rather than partitioned into discrete clusters. By applying PCA to the population’s RM maps, we uncovered that the most prominent latent structure (PC1) was unipolar across the cortex and strongly correlated with the traditional whole-brain “age gap” metric, reflecting a global shift in maturational timing. In contrast, higher-order components exhibited bipolar distributions, capturing coordinated developmental asynchrony in opposing directions across different brain networks. These axes of developmental imbalance aligned closely with FC gradients. This alignment suggests that developmental imbalances are constrained by the brain’s intrinsic functional hierarchy, rather than emerging as spatially arbitrary deviations.

Although developmental imbalance is hypothesized as a main neural mechanism of psychopathological problems [8, 14, 34, 53], empirical evidence is still lacking. Here, we provided the most direct neurobehavioral evidence to date that an individual’s specific spatial pattern of developmental imbalance significantly and reliably predicts their unique psychopathological profile. Recent studies have established the S-A axis as a principal gradient of both cortical organization and normative development [14–16, 54]. We further revealed that the critical imbalance pattern that predicts psychopathological risks aligned with the S-A axis, capturing the tension between the brain’s top-down regulatory functions and bottom-up reward-seeking in shaping psychopathological risks. In particular, our findings suggest accelerated maturation of the higher-order association systems relative to sensorimotor regions may enhance social and reward-oriented cognitive processes, increasing sensitivity to peer contexts and the perceived benefits of risk behaviors such as alcohol use. In contrast, when the association cortex matures more slowly than the sensorimotor circuits, individuals may experience stronger bottom-up drives but weaker top-down inhibitory control, resulting in increased impulsivity, difficulties in attention regulation, and internalizing symptoms. These findings establish the S-A axis as a continuous “liability spectrum,” where the relative maturational timing of these distinct functional systems dictates an individual’s specific style of psychiatric vulnerability. Thus, our study suggests that these system-level patterns of developmental imbalance, rather than developmental deviation in isolated regions, might be a fundamental driver for generic psychopathological vulnerability.

### Limitations

Several limitations should be considered when interpreting these results. First, our analyses relied on a group-level atlas. While necessary for cross-subject correspondence, this approach neglects the individual variability of functional topography (e.g., the specific layout of network boundaries) [55, 56]. This “functional misalignment” may attenuate the precision of our maturity estimates, particularly in association cortices where inter-subject variability is highest [56]. Future iterations could leverage precision functional mapping techniques to define individual-specific parcels [57, 58], potentially enhancing the sensitivity of the Relative Maturity index. Second, while we utilized large-scale datasets, the predictive fidelity of region-wise models (average r approx. 0.4) is low. This limits the confidence of our inferences. Predicting regional maturity with higher fidelity will likely require larger datasets with significantly longer acquisition times per subject [59]. Third, our analyses are restricted to the cerebral cortex; future work should extend this framework to include subcortical structures. Lastly, models trained on the whole cohort might lack sensitivity in highlighting sex differences in relative maturity. We observed patterns consistent with prior literature [1], with some regions showing higher RM in females during earlier developmental stages and lower RM at later stages (Supplementary Fig. 13). However, these sex differences were modest and did not survive multiple-comparison correction, thus unlikely influencing our main findings. A more targeted investigation of sex-specific developmental trajectories is warranted.

---

**Supplementary Figure 1.**
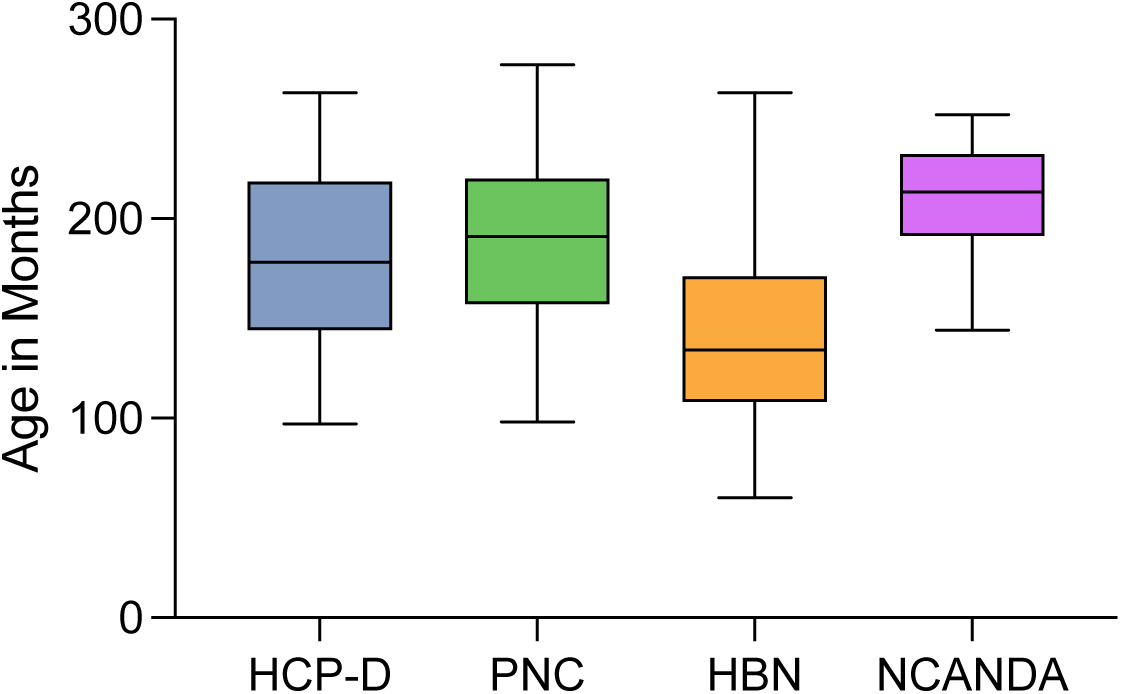
Age distribution of the datasets used in this study. Participant age distributions (in months) are shown across the four independent cohorts: HCP-D, PNC, HBN, and NCANDA. The three cross-sectional cohorts (HCP-D, PNC and HBN) have similar age range, while the longitudinal cohort (NCANDA) has relatively high age.

**Supplementary Figure 2.**
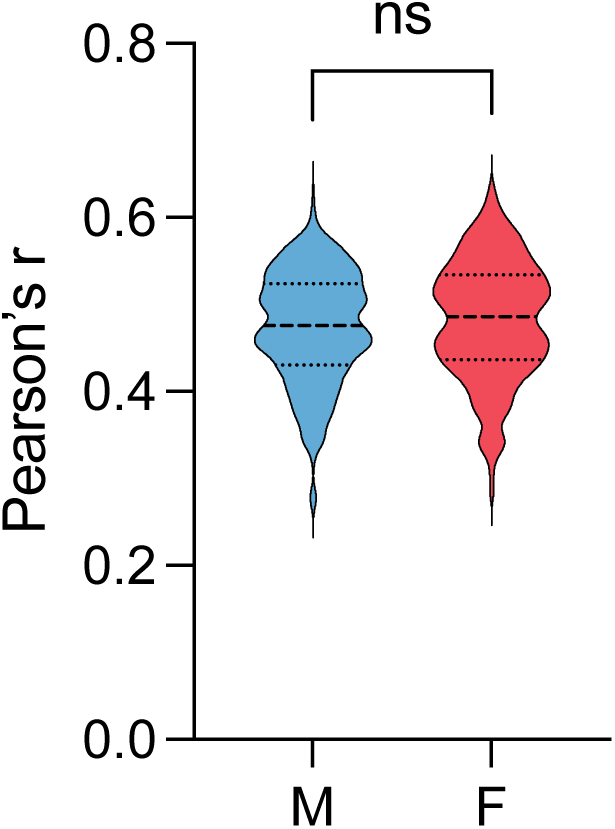
Sex difference in age prediction accuracy. In the HCP-D cohort (the cohort with highest prediction accuracy according to Fig. 2), we trained the KRR models on a subset of sex-balanced samples (N=158) and tested the models on an independent set of 158 males and 158 females. The experiment was repeated 20 times using different random seed for data split. The resulting prediction accuracy showed no significant sex differences in accuracy.

**Supplementary Figure 3.**
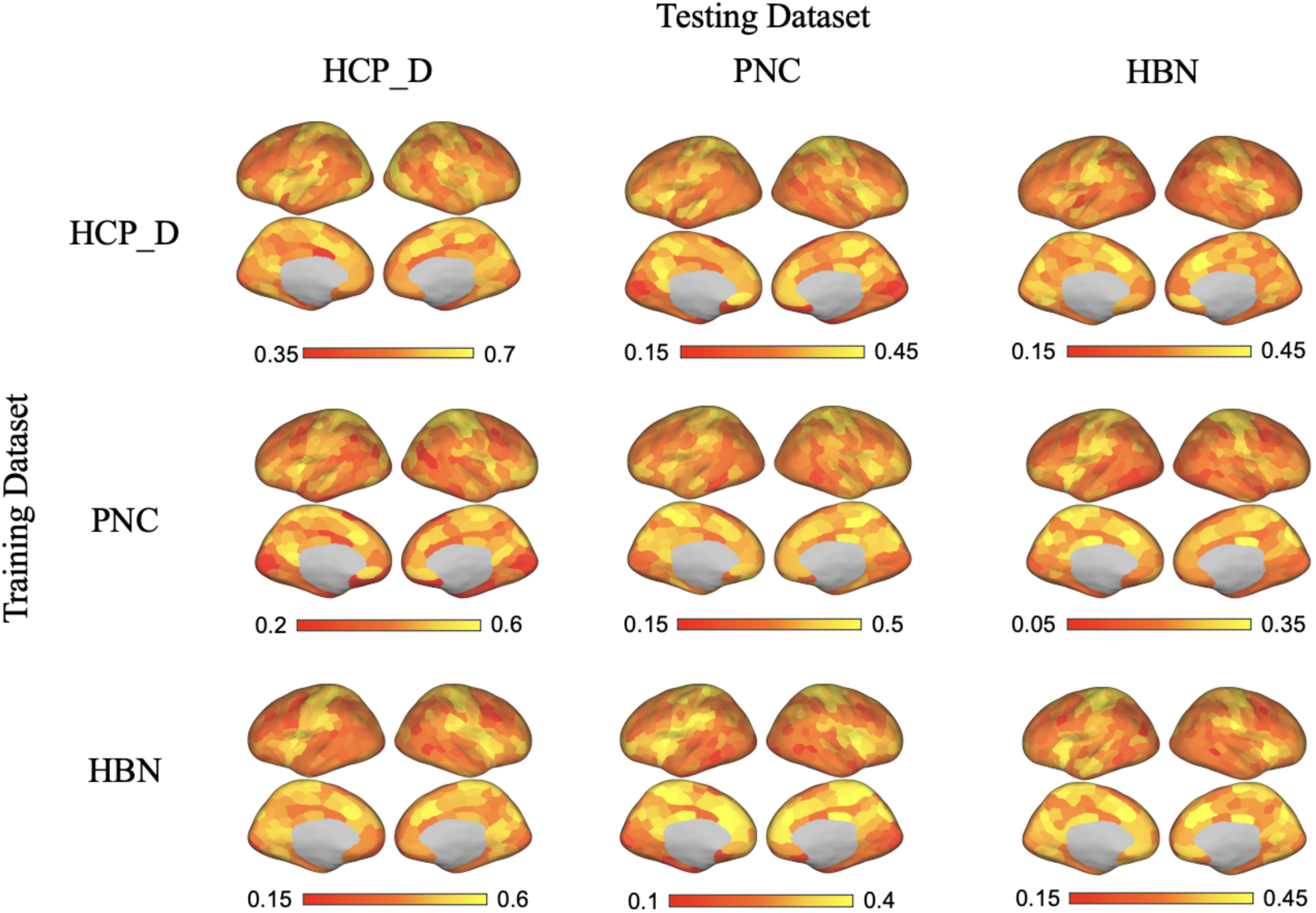
Spatial topography of regional age prediction accuracy. Brain maps display the prediction accuracy of regional machine-learning models across various training and testing cohorts. Rows indicate the dataset utilized to train the models (HCP-D, PNC, or HBN), while columns denote the dataset used for testing. Maps on the diagonal represent within-dataset model performance (evaluated by 10-fold cross-validation). In contrast, maps on the off-diagonal represent out-of-sample, cross-cohort generalization, demonstrating the spatial consistency of developmental signatures across independent datasets. Color bars denote the specific range of prediction accuracy (Pearson’s r) for the train-test pair.

**Supplementary Figure 4.**
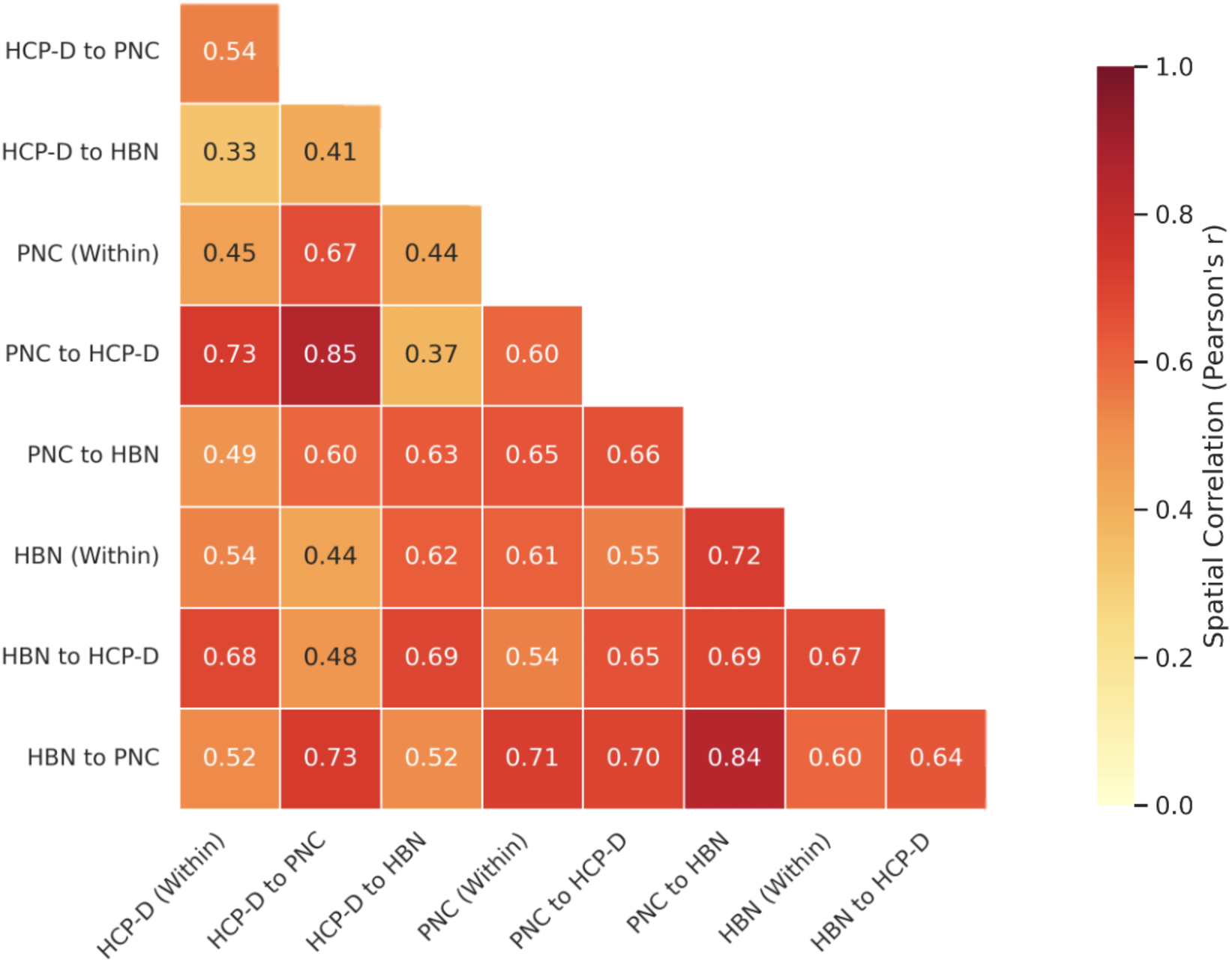
Spatial correlation of regional prediction accuracy across training and testing settings. The lower-triangular heatmap displays the pairwise spatial similarity (Pearson’s r) between the regional age prediction accuracy maps derived from various training-testing dataset combinations (HCP-D, PNC, and HBN). Axis labels indicate the specific training-testing setting (e.g., “HCP-D to PNC” denotes models trained on HCP-D and tested on PNC, whereas “Within” denotes 10-fold cross-validation within the same cohort). The consistent moderate-to-high spatial correlations confirm that the spatial topography of regional age prediction accuracy is highly reproducible.

**Supplementary Figure 5.**
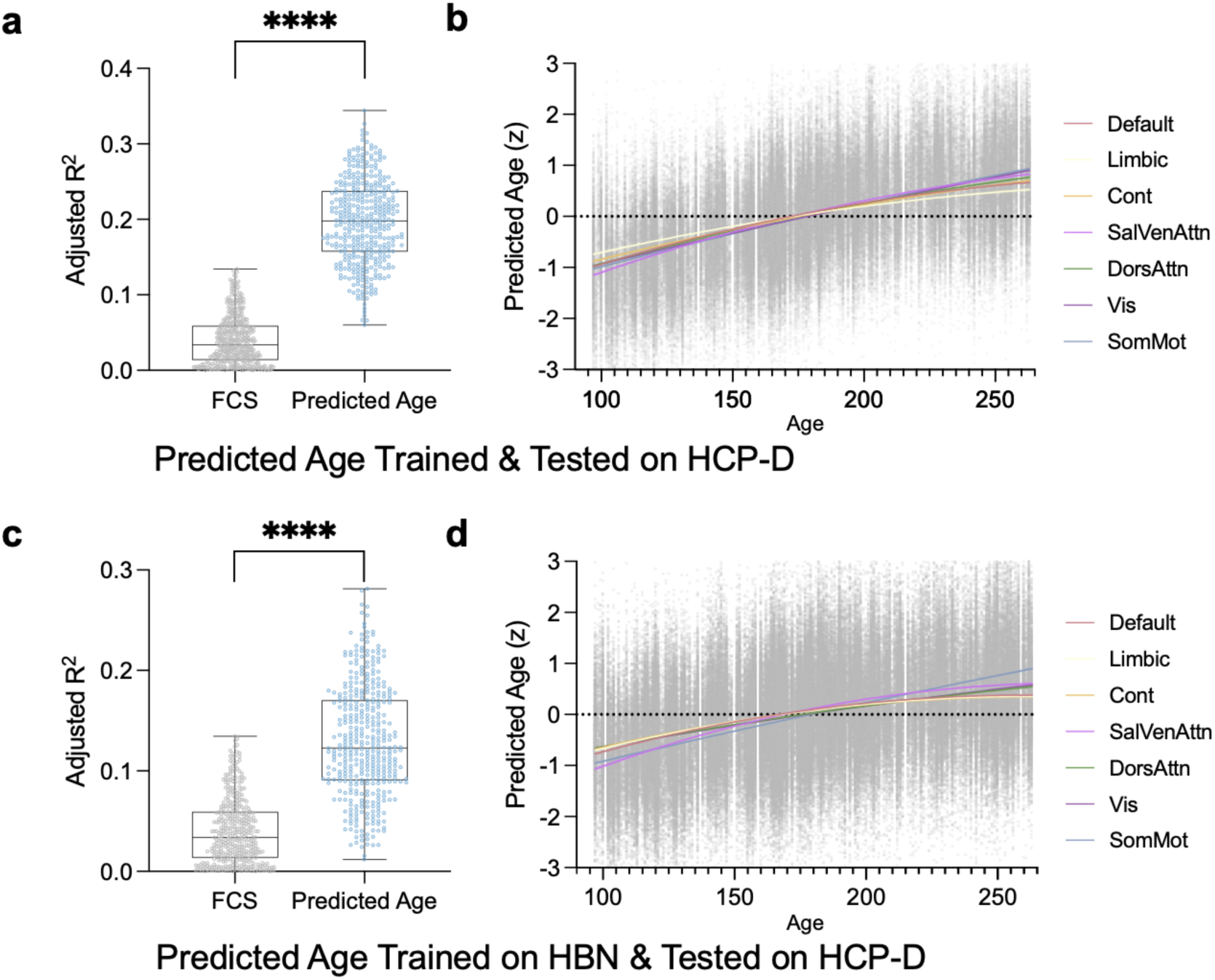
Validation of sensitivity to developmental changes of FCS and predicted age. Evaluation of the sensitivity to developmental changes of regional age prediction is conducted on the HCP-D cohort. Models are either trained within HCP-D using 10-fold cross validation (Top row) or trained on an independent dataset of HBN (Bottom row). **a, c**: Pairwise comparison of the age-related variance explained (Adjusted R²) by Mean Functional Connectivity Strength (FCS) and predicted age across all brain regions. The predicted age explains significantly higher variance than FCS (**** indicates p < 0.0001, paired t-test). **b, d**: Network-level developmental trajectories of the regional age prediction. Scatter plots display individual region-level FCS or predicted age (standardized to z-scores) as a function of age (in months). Colored curves illustrate ordinary least squares (OLS) regression fits of 7 functional networks.

**Supplementary Figure 6.**
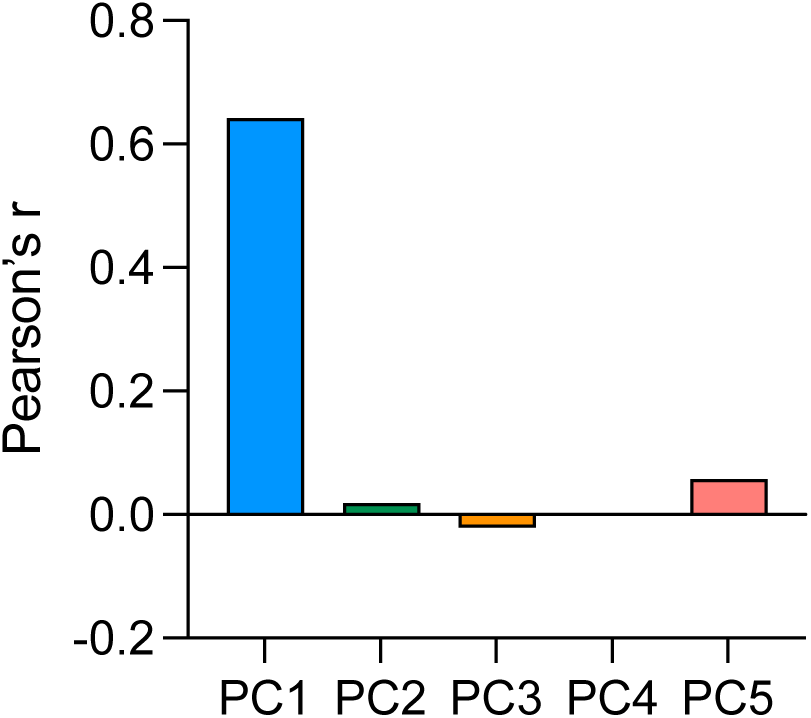
Correlation between principal components of regional relative maturity maps and whole-brain relative maturity. The bar plot displays the Pearson correlation between the first five principal components (PCs) of the regional relative maturity (RM) maps and the whole-brain relative maturity, derived by a global model that predicts age from whole-brain functional connectivity. PC1 exhibits a strong positive correlation, confirming it captures a uniform, brain-wide maturation trend. Higher-order components (PC2–PC5) show near-zero correlations.

**Supplementary Figure 7.**
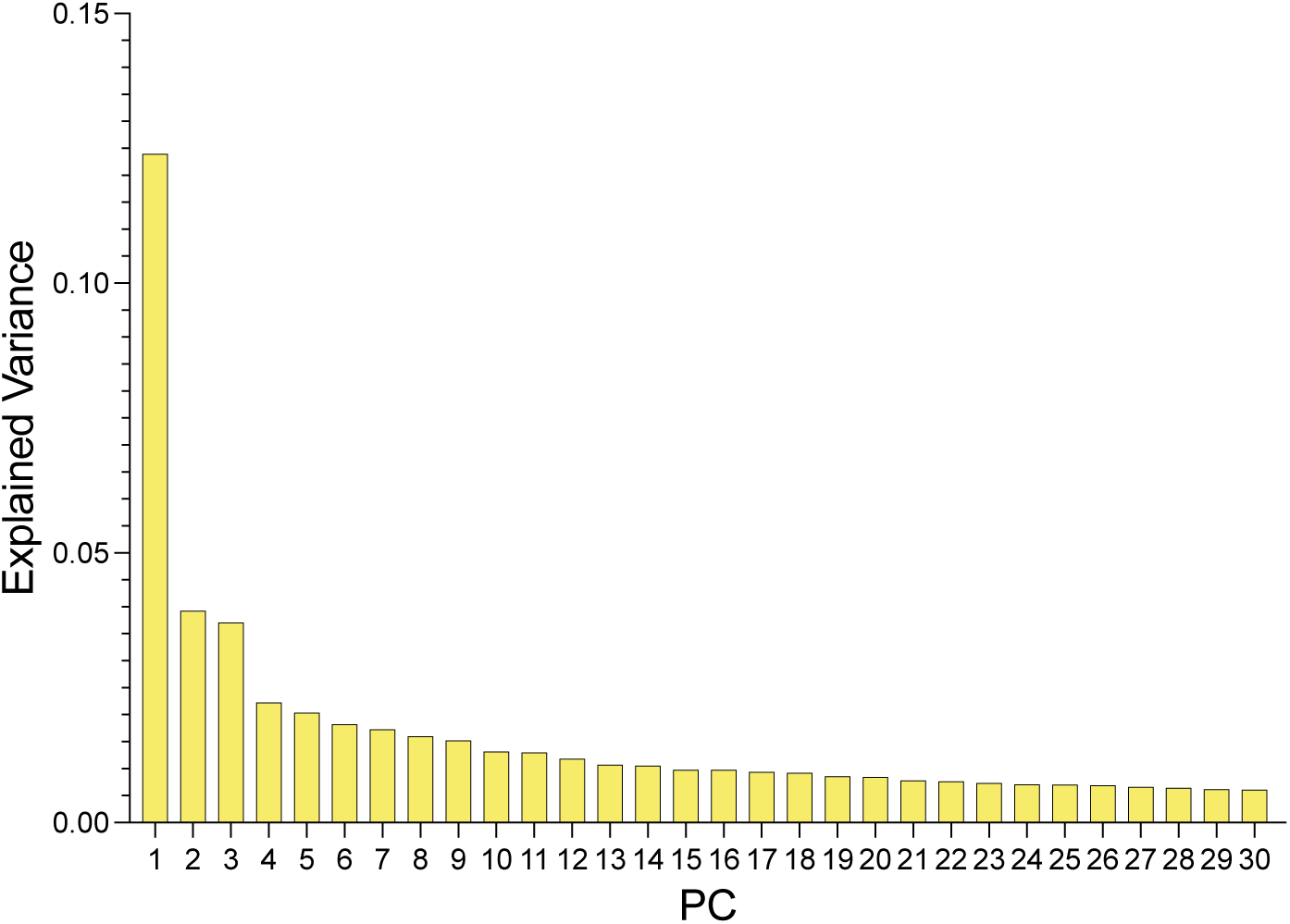
Explained variance of principal components in the combined cross-sectional cohort. The bar plot illustrates the explained variance for the top 30 PCs derived from the combined cross-sectional datasets (HCP-D, PNC and HBN).

**Supplementary Figure 8.**
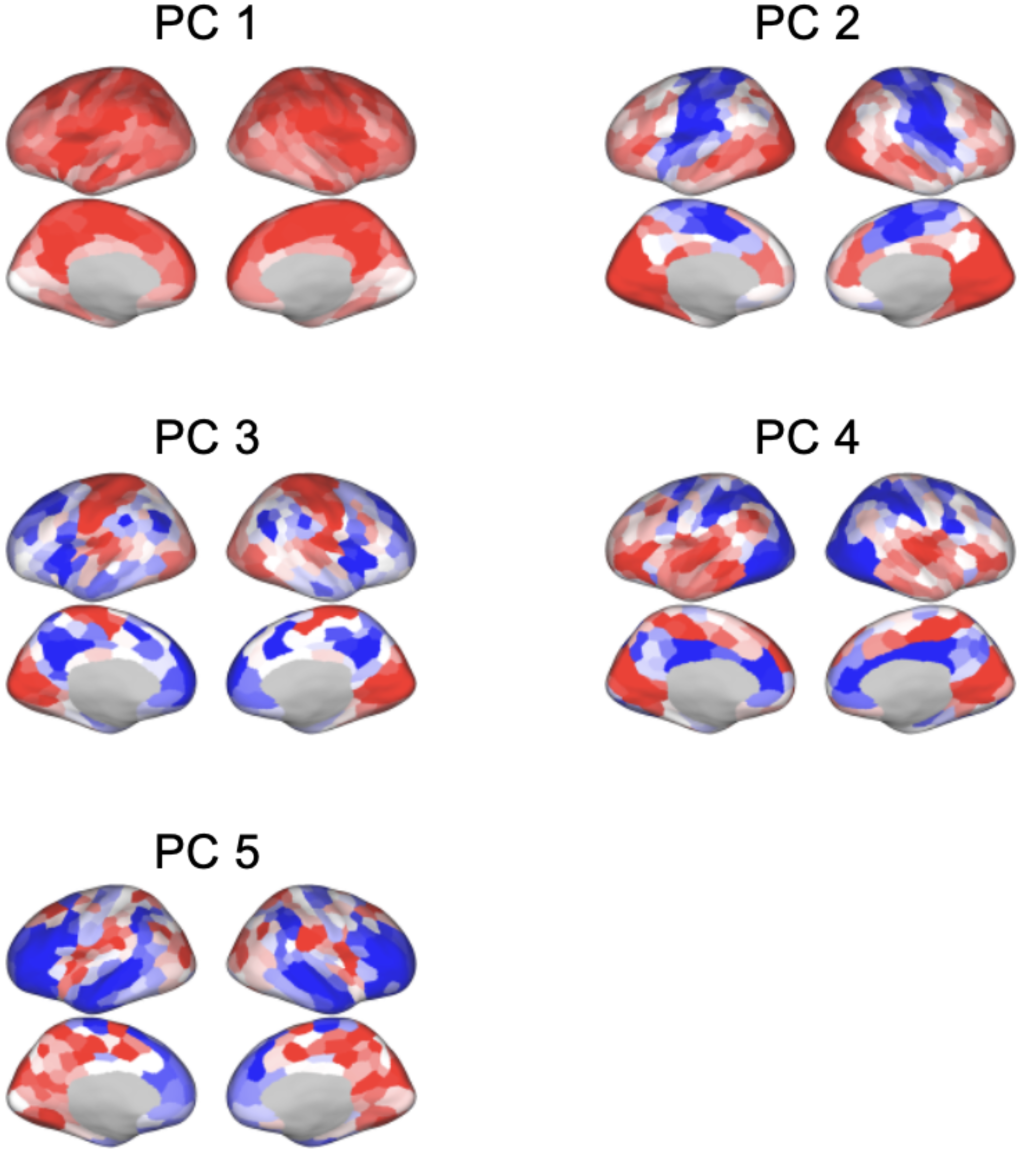
Replication of latent spatial structure of relative maturity in the NCANDA cohort. Brain maps display the loadings of the first five PCs derived from the individualized RM maps within the longitudinal NCANDA dataset. Similar to our results derived from the combined cross-sectional cohort (HCP-D, PNC and HBN, see Fig. 5), PC1 captures a unipolar, global maturation trend across the cortex, while higher-order components (PC2–PC5) exhibit bipolar spatial distributions, reflecting the latent structure of developmental imbalances.

**Supplementary Figure 9.**
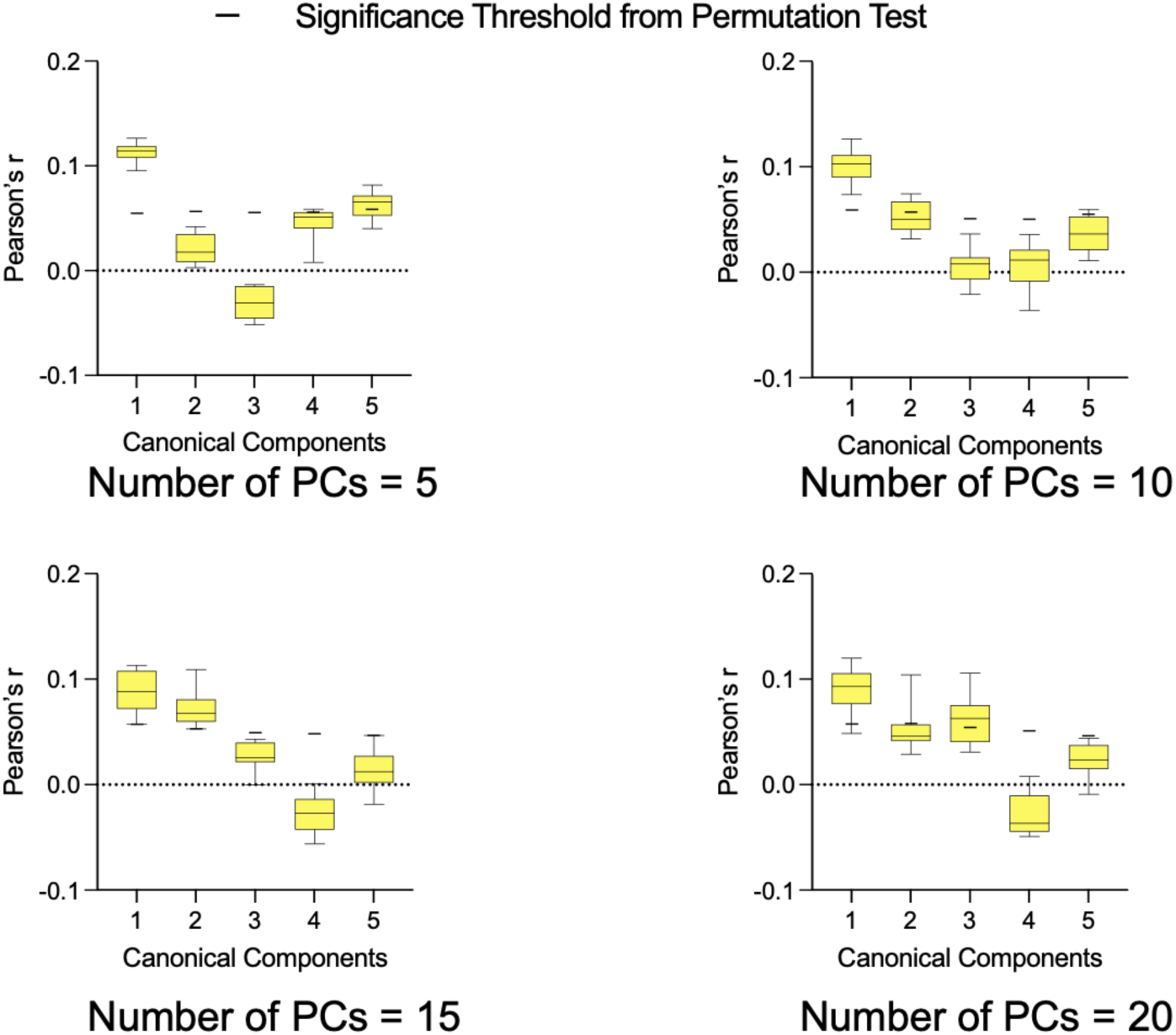
Out-of-sample prediction accuracy of canonical components. The boxplots show the out-of-sample cross-validation accuracy (Pearson’s r) for the first five canonical components linking latent RM modes to psychopathological dimensions. To evaluate the stability of the brain-behavior association, the Canonical Correlation Analysis (CCA) was evaluated across varying dimensionalities of the input data, using either 5, 10, 15, or 20 PCs of the RM maps. The short horizontal lines denote the component-specific significance thresholds derived from permutation test (p=0.05). Across all evaluated PC thresholds, the first canonical component consistently achieves statistically significant out-of-sample prediction.

**Supplementary Figure 10.**
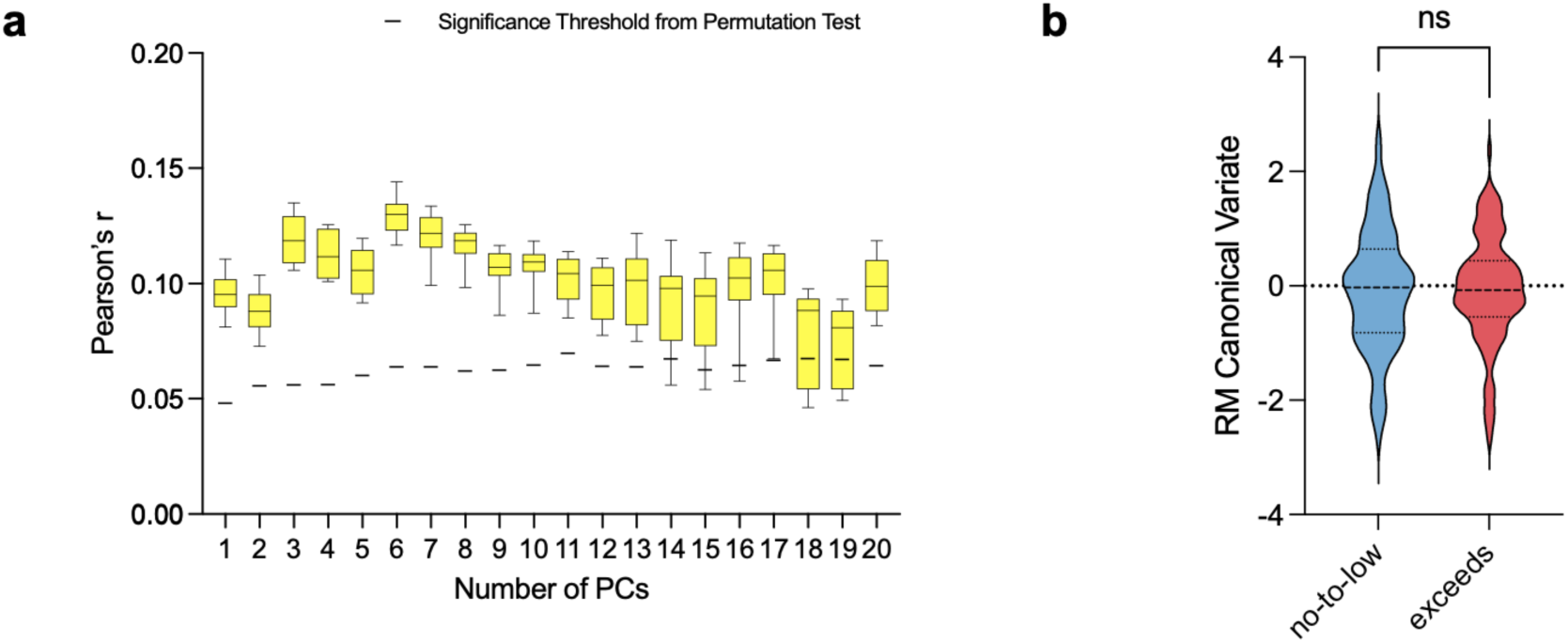
Robustness of the brain-behavior association to baseline alcohol use. **a,** Out-of-sample prediction accuracy (Pearson’s r) of the first canonical component is evaluated using 10-fold cross-validation in 645 NCANDA participants with no-to-low alcohol use at baseline. To ensure the brain-behavior associations were not driven by baseline heavy drinking, we systematically varied the number of input PCs of the RM maps from 1 to 20 in CCA. The boxplots display the distribution of out-of-sample prediction accuracy of the first canonical component across 10 random seeds for training-testing data split. Horizontal lines indicate the significance thresholds established by permutation testing (p=0.05). Regardless of the choice of number of PCs, the first canonical component demonstrates robust, statistically significant out-of-sample correlation that peaks when utilizing 6 input PCs. **b,** Comparison of the RM canonical variate between participants who exceeded the binge drinking threshold at baseline and a matched subset drawn from the no-to-low alcohol use at baseline group. The two groups were matched by age, sex, and sample size. No significant difference was observed, confirming that the canonical variate scores are not artificially driven by heavy baseline drinking.

**Supplementary Figure 11.**
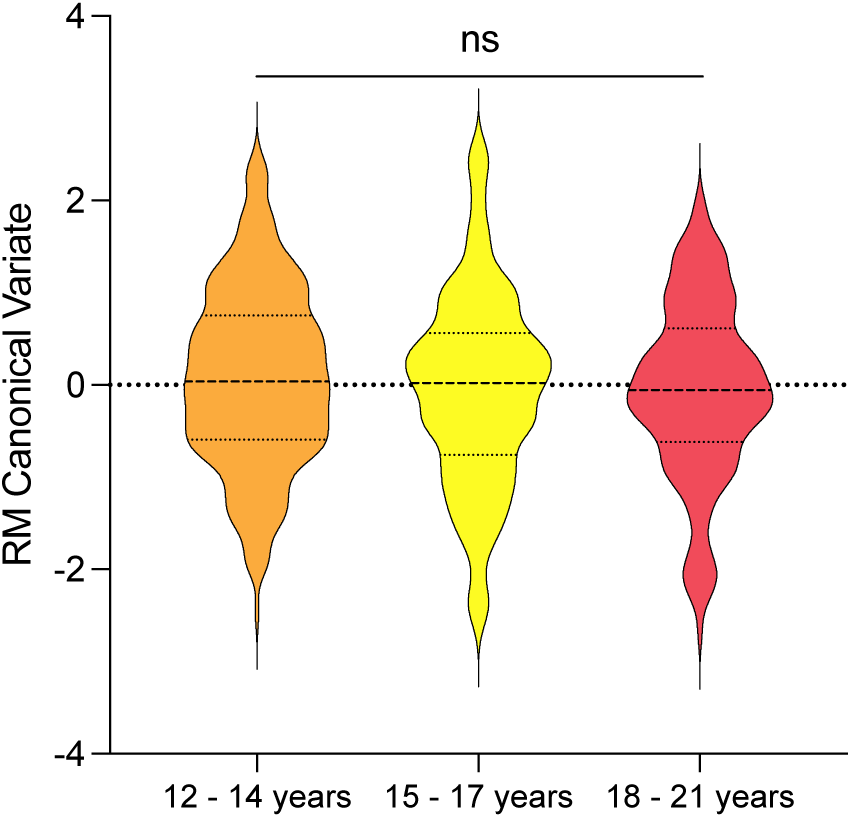
Comparison of the RM canonical variate across baseline age bands. The violin plots show the distribution of the RM canonical variate for subjects in three baseline age ranges: 12-14, 15-17, and 18-21 years (144–180, 180–216, and 216–252 months). A one-way ANOVA and three pairwise unpaired t-tests confirmed there are no statistically significant differences between any of the age bands. This establishes that the RM canonical variate is not confounded by baseline age effects.

**Supplementary Figure 12.**
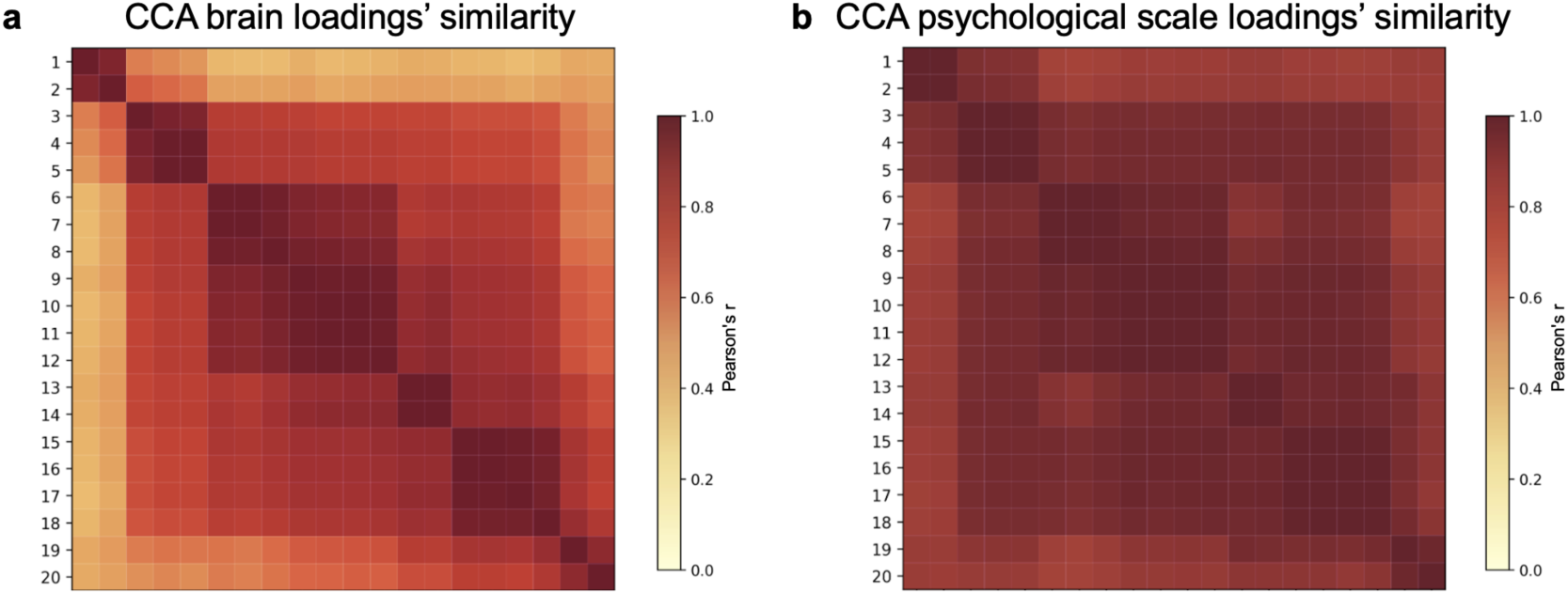
Stability of CCA brain and psychological scale loadings across varying input dimensionalities. Pairwise similarity matrices compare the loadings of the first canonical component across CCA models with different numbers of PCs of RM maps as the input. **a,** Pearson’s correlation between the brain loading maps. **b,** Pearson’s correlation between the psychological scale loadings. For both panels, the axes denote the number of PCs (ranging from 1 to 20) utilized as brain features to fit the CCA models. The consistently high correlations, particularly across models using 3 to 18 PCs (Pearson’s r > 0.75), demonstrate that the association between developmental imbalance and psychopathological scales is highly stable and not just an artifact of the chosen input dimensionality.

**Supplementary Figure 13.**
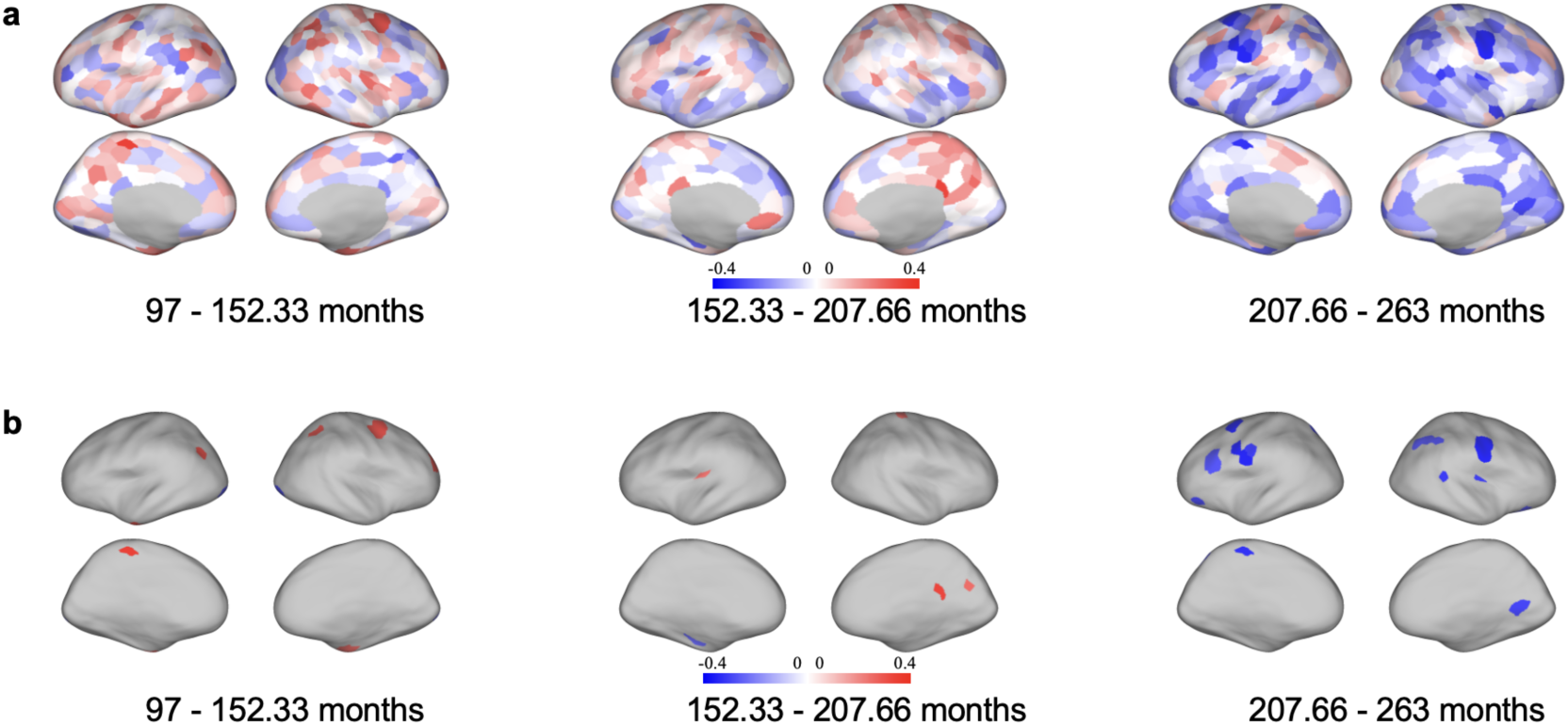
Evaluation of sex differences in regional RM across development. RM of the HCP-D cohort is derived using within-dataset cross-validation to assess potential sex differences in maturational timing. Subjects were stratified into three equal age bands spanning 97 to 263 months. **a,** Brain maps show the mean RM differences between sexes (Female mean minus Male mean). Red indicates regions where females exhibit higher RM, whereas blue indicates higher RM in males. **b,** Brain maps show the regions with significant sex differences (uncorrected p < 0.05, two-sample t-test). While a sparse number of regions demonstrate advanced maturation in females during the earliest developmental window, no regions survive multiple comparison correction (FDR corrected p > 0.05). These findings indicate that there exist modest interpretable sex differences in maturational timing, yet not pronounced enough when the age prediction models are trained on the whole cohort.

**Supplementary Figure 14.**
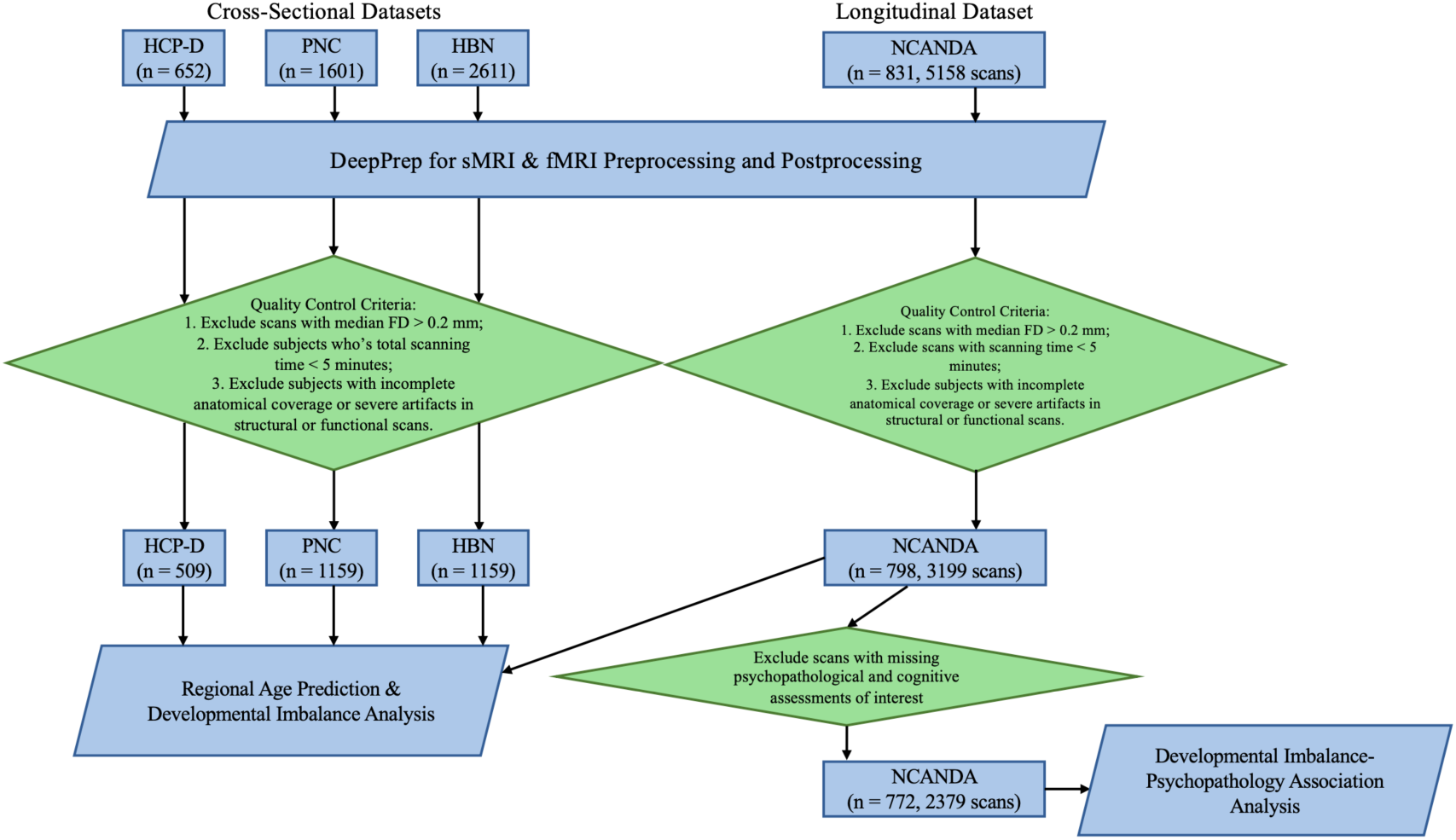
Data processing and exclusion pipeline. An overview of our data processing and quality control (QC) pipeline for the cross-sectional (HCP-D, PNC, HBN) and longitudinal (NCANDA) cohorts. We leveraged DeepPrep for structure and functional MRI processing. Subsequently, rigorous QC criteria were applied to exclude scans exhibiting excessive head motion (median framewise displacement >0.2 mm), insufficient scan duration (<5 min), incomplete anatomical coverage or severe imaging artifacts. NCANDA participants with structural anomaly at baseline visits were excluded. For the developmental imbalance-psychopathology association analyses within the NCANDA cohort, we further excluded visits lacking a complete battery of psychopathological assessments.

**Supplementary Table 1.**
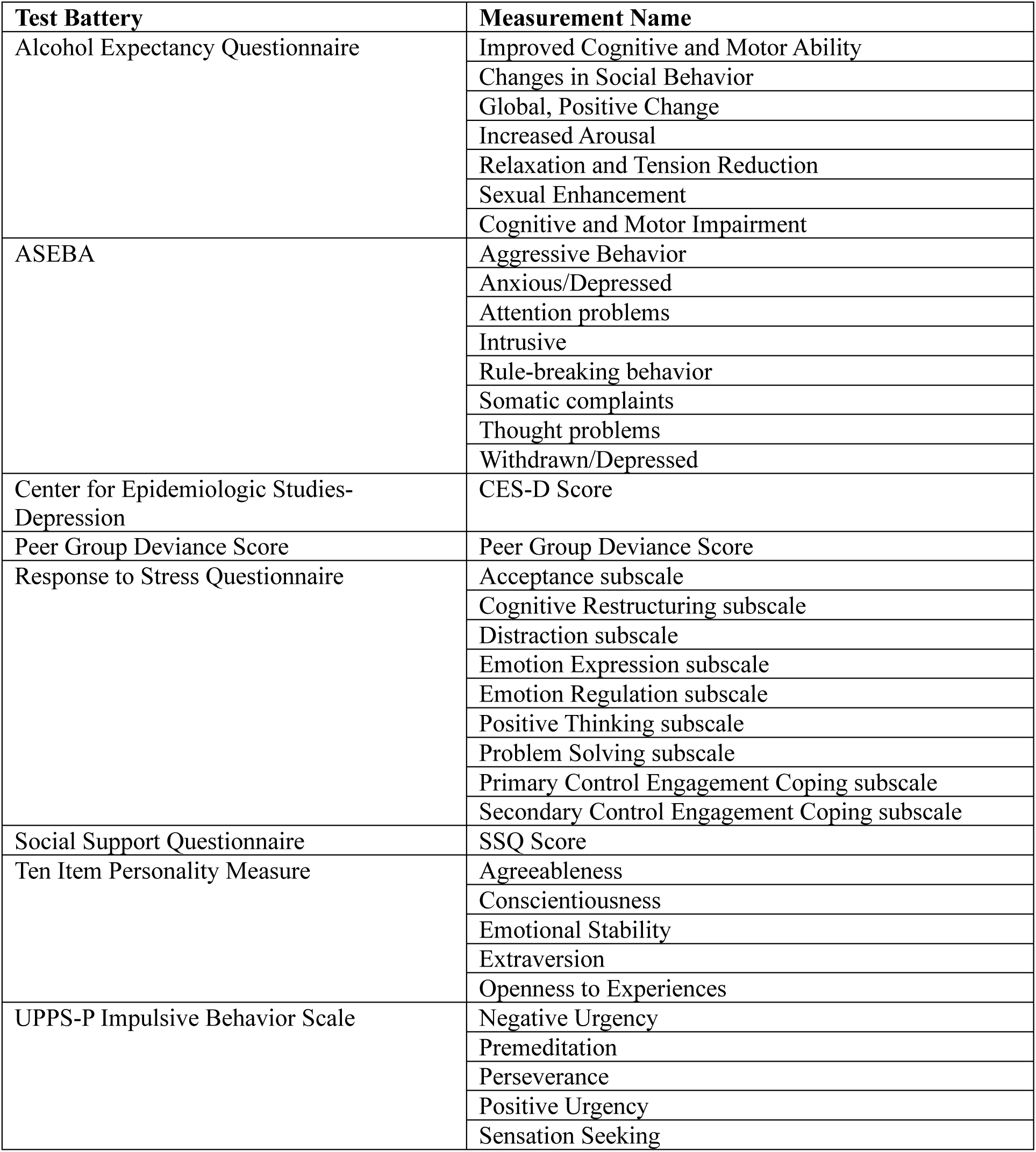
Psychopathological Measurements included in NCANDA analysis.

## Methods

### Participants

This study leveraged four large-scale, independent datasets. The three cross-sectional cohorts were the Human Connectome Project Development (HCP-D; *n* = 509, 8-21 years, 271 females), the Philadelphia Neurodevelopmental Cohort (PNC; *n* = 1,159, age 8-23 years, 623 females), and the Healthy Brain Network (HBN; *n* = 1,159, age 5-21 years, 472 females). Longitudinal validation was performed using the National Consortium on Alcohol and Neurodevelopment in Adolescence (NCANDA; *n* = 798, 12–22 years, 390 females, 4.01±1.98 longitudinal visits per participant, 3,199 visits in total). Participants were selected based on the following preprocessing steps (see Supplementary Fig. 14).

### Quality Control and Data Preprocessing

Across all cohorts, participants were excluded if they met any of the following conditions: (1) high head motion, defined as median Frame Displacement > 0.2 mm; (2) insufficient data quantity, defined as total scanning time < 5 minutes; or (3) incomplete anatomical coverage or severe artifacts in structural or functional scans.

Structural and functional MRI data were preprocessed using the DeepPrep [60-62] pipeline. Functional MRI data underwent the following specific processing steps: (1) bandpass filtering (0.01–0.08 Hz); (2) regression of confounds, including head motion parameters and global signal derived from DeepPrep; (3) smoothing using a 6-mm full-width at half-maximum kernel; and (4) projection to the fsaverage6 surface space (40,962 vertices) [63]. Finally, regional time series were extracted using the Schaefer 400-parcel atlas (Kong’s version [64]) to compute FC matrices for subsequent analysis.

### Regional Age Prediction

For region-specific age prediction, we employed a Kernel Ridge Regression framework, which demonstrated superior performance in our preliminary validation [38]. Prior to model training, we performed a covariate regression procedure to isolate age-related effects from potential confounds. Specifically, we fitted a linear regression model on the training set to remove the confounding effects of sex and mean head motion (Frame Displacement) from each FC feature. The coefficients derived from the training set were then applied to correct the features in the testing set, ensuring no data leakage. These corrected FC profiles served as the input features for the region-specific Kernel Ridge Regression models.

After training a region-specific prediction model, we applied it to a testing population and derived their predicted age as the maturity index of that region. To compare the predicted age and FCS, both metrics were first standardized by *z*-scoring across all testing subjects and regions to account for differences in scale and distribution. Then for each region, we evaluated the metrics’ sensitivity to developmental changes using OLS regression:

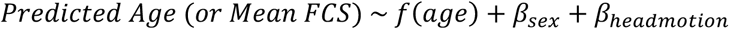

Here, head motion is estimated by the mean Frame Displacement during fMRI scan. For either metric, we calculated the unique variance explained by age as the difference in adjusted *R*^2^ between a full model (including age) and a reduced model (covariates only). In addition, to assess whether a metric captured network-level age effects, we aggregated the metrics of all regions within each of Yeo’s 7 intrinsic functional networks and fitted network-level OLS models.

Lastly, we derived the region’s RM by first fitting a linear regression model to regress chronological age out of the predicted brain age. The resulting residuals were then z-scored. This procedure was also used to define global relative maturity by training and testing a separate Kernel Ridge Regression model to predict age prediction from the whole brain FC.

### Association between Developmental Imbalance and Psychopathology in the NCANDA Cohort

#### Psychopathological and Behavioral Assessments

To capture a comprehensive profile of adolescent psychiatric vulnerability, we followed our prior work [42] and identified 37 out of the 45 socioemotional scores from the NCANDA study. Note, we removed the life experiences measures and Karolinska Sleepiness Scale used in [42] to only focus on psychopathological symptoms. These measures span 7 domains: Childhood Behavioral Checklist, UPPS-P Impulsivity, general depression severity, Ten-Item Personality Inventory, Responses to Stress Questionnaire, Alcohol Outcome Expectancies, and Social and Environmental Factors. The full list of psychopathological measures is given in Supplementary Table 1. From the 798 NCANDA participants with usable rs-fMRI scans during ages 12-22, 2,369 longitudinal visits of 772 participants had complete data on all 37 psychopathological scales.

#### Canonical Correlation Analysis

To investigate the clinical relevance of developmental imbalance, we first extracted RM maps of the NCANDA cohort. Specifically, we applied the regional age-prediction models trained on the combined cohort of three cross-sectional datasets (HCP-D, PNC, and HBN) to predict age for each NCANDA scan. We then derived individualized Relative Maturity (RM) maps and performed PCA on the maps. We employed CCA to quantify the high-dimensional covariate patterns between the PC scores of the RM maps and the 37 psychopathological scales. Specifically, CCA derives a set of canonical components, each consisting of a pair of canonical variates: one representing a linear transformation of the PC scores and the other representing a linear transformation of the psychopathology scales. CCA aims to maximize the correlation between the two canonical variates. The resulting canonical correlation value reflects the overall strength of the mapping between developmental imbalance and psychopathology.

#### Model Validation and Interpretation

To robustly estimate the out-of-sample generalizability of the canonical components, we implemented a subject-level 10-fold cross-validation, where all longitudinal scans from a given participant were assigned to the same fold. The out-of-sample correlation between the two transformed canonical variates was computed on the testing fold and averaged across all 10 folds. To ensure stability, this stratified procedure was repeated using 10 distinct random seeds, and the statistical significance of the out-of-sample canonical correlation was determined via permutation testing (1000 permutations).

To identify which specific psychopathological scales drove the association in a canonical component, we computed the canonical loading of each scale, i.e., the Pearson correlation between the original psychopathological scale and the corresponding canonical variate. Similarly, we derived canonical loadings of the brain PC scores. To interpret these loadings neurobiologically, we back-projected them into the original 400-region cortical space. Since each PC corresponds to a spatial weight map across the cortex, we computed the weighted sum of these PC spatial maps, scaled by their respective canonical loadings. This back-projection yielded a regional brain loading map, visualizing the specific directional contribution of each cortical region’s relative maturity to the significant brain-behavior canonical variate.

## Data Availability

The Philadelphia Neurodevelopmental Cohort and the Healthy Brain Network datasets are publicly available in the RBC website (https://reprobrainchart.github.io/), the Human Connectome Project-Development are available in the NIMH Data Archive (https://nda.nih.gov/). The National Consortium on Alcohol and Neurodevelopment in Adolescence is available at http://www.ncanda.org/datasharing.php.

## Code Availability

Code is available upon request.

## Acknowledgment

This work was supported by National Institutes of Health grants U24 AA021697 (K.M.P.), R01 DA057567 (K.M.P.), and R00 AA028840 (Q.Z.); the Brain & Behavior Research Foundation (BBRF) Young Investigator Award (Q.Z.); the National AI Research Resource (NAIRR) Pilot Resource Grant (Q.Z.); Bowers Foundation grant through the Ann S. Bowers Women’s Brain Health Initiative (A.K.)

## Author Contributions

Conception: Q.H. and Q.Z. Design: Q.H., L.M., B.J., K.M.P, A.K. and Q.Z. Data acquisition, analysis and interpretation: Q.H. and Q.Z. Manuscript writing and revision: Q.H., L.M., B.J., K.M.P, A.K. and Q.Z. All authors approved the final manuscript.

## Competing interests

The authors declare no competing interests.

## Notes

### Competing Interest Statement

The authors have declared no competing interest.

